# A noncanonical Pol III-dependent, Microprocessor-independent biogenesis pathway generates a germline enriched miRNA family

**DOI:** 10.1101/2025.04.11.648421

**Authors:** Rima M. Sakhawala, Reyhaneh Tirgar, Karl-Frédéric Vieux, Dustin Haskell, Guoyun Yu, Anna Zinovyeva, Katherine McJunkin

## Abstract

MicroRNAs (miRNAs) are short RNAs that post-transcriptionally regulate gene expression. In canonical miRNA biogenesis, primary miRNAs are transcribed from intergenic loci or intronic regions by RNA polymerase II, sequentially cleaved by the Microprocessor complex and Dicer, and resulting mature miRNAs are loaded into Argonaute to repress target mRNAs. A minority of miRNAs are generated via noncanonical biogenesis pathways that bypass the Microprocessor and/or Dicer. Here, we describe a new Pol III-dependent, Microprocessor -independent, and Dicer-dependent biogenesis pathway exemplified by the *mir-1829* family in *C. elegans*. Although the *mir-1829* family loci reside in intronic regions of protein-coding genes, we show that the miRNAs are derived from independent Pol III transcripts. Unlike other Pol III-dependent miRNAs, the *mir-1829* family small RNAs are the dominant species derived from their loci, rather than fragments of a larger functional noncoding RNA. These germline-enriched miRNAs are loaded in multiple miRNA Argonautes, including the recently-characterized germline Argonaute ALG-5, which we demonstrate is repressive when tethered to a reporter transcript. We extend these findings, identifying additional Pol III-transcribed and noncanonical small RNAs in *C. elegans* and human datasets, including human miR-4521. These young, noncanonical miRNAs may represent an early snapshot in the evolution of de novo miRNA genes.

## Introduction

MicroRNAs (miRNAs) are small, ∼22-nucleotide regulatory RNAs that selectively repress gene expression during development and differentiation. In canonical miRNA biogenesis, primary miRNAs are transcribed from intergenic loci or intronic regions by RNA polymerase II (Ha and Kim 2014). These transcripts are then processed by the Microprocessor (MP), a complex that consists of the catalytic RNase III enzyme Drosha and a homodimer of the RNA-binding protein DGCR8 (known as DRSH-1 and PASH-1 in *C. elegans*, respectively) (Lee et al. 2003; Denli et al. 2004; Gregory et al. 2004; Han et al. 2004). The MP cleaves the stem of hairpin structures of primary miRNAs (pri-miRNAs) to produce miRNA precursors, which are further processed into small RNA duplexes by the RNase III enzyme Dicer (DCR-1 in *C. elegans*) (Hutvágner et al. 2001; Knight and Bass 2001; Grishok et al. 2001). One strand of the resulting duplex is preferentially loaded into an Argonaute (Ago) protein, forming the miRNA-Induced Silencing Complex (miRISC), which can target mRNAs via base pairing to their 3’ UTR to repress translation and/or mediate the recruitment of factors that cause mRNA decay (Gebert and MacRae 2019).

Beyond canonical miRNA biogenesis, multiple alternative biogenesis pathways bypass the MP and/or Dicer. One of the best-studied examples is mirtron biogenesis, in which a short intron takes on a hairpin-like structure after debranching; this hairpin is then processed as a miRNA precursor by Dicer, thus bypassing the requirement for MP (Berezikov et al. 2007; Okamura et al. 2007; Ruby et al. 2007). Some mirtron-like molecules, termed “Agotrons” bypass both MP and Dicer and associate with Agos as full-length introns (Hansen et al. 2016). Short, 5’ 7-methylguanine (m^7^G) capped Pol II transcripts can also bypass the MP, being directly processed by Dicer before Ago loading (Sheng et al. 2018; Babiarz et al. 2008; Xie et al. 2013; Zamudio et al. 2014). Dicer-independent biogenesis is exemplified by vertebrate miR-451, which is cleaved by MP, but then relies on the catalytic activity of Ago to further its maturation (Cheloufi et al., 2010; Cifuentes et al., 2010; S. Yang et al., 2010).

In addition to these Pol II transcripts that undergo noncanonical processing, Pol III-derived transcripts can also be fragmented – generally in a MP-independent, Dicer-dependent manner – giving rise to Ago-loaded small RNAs. Regions of tRNAs that form hairpins can be further processed by Dicer to produce miRNA-like small RNAs (Babiarz et al. 2008; Kuscu et al. 2018; Maute et al. 2013; Martinez et al. 2017). Similarly, specific snoRNAs and the <u>s</u>mall <u>N</u>F90 <u>a</u>ssociated <u>R</u>NA <u>A</u> (snaR-A) serve as sources for distinct Dicer-dependent small RNA species (Ender et al. 2008; Lemus-Diaz et al. 2020; Stribling et al. 2021). Dicer can also generate small RNAs from hairpins formed by transcribed SINE elements (Babiarz et al. 2008). Overall, diverse Pol III transcripts give rise to less abundant small RNA products that can function as miRNAs.

In this study, we present a new noncanonical miRNA biogenesis pathway exemplified by the *mir-1829* family in *C. elegans*. Although they reside within long introns of Pol II-derived host genes, the *mir-1829* family members are derived from independent transcriptional units transcribed by Pol III. The *mir-1829* family is MP-independent but Dicer-dependent. The *mir-1829* family is enriched in the germline and, in particular, enriched in ALG-5, a germline specific Ago protein; we show via a tethering assay that ALG-5 is repressive, and thus likely functions similarly to other miRNA Agos. Unlike previously-identified Pol III-dependent miRNAs, the *mir-1829* family small RNAs appear to be the primary product of these loci, not derivatives of a more abundant functional species. We further expanded our search for additional noncanonical and Pol III-derived miRNAs by cross-referencing datasets in which MP, Dicer, and Pol III are depleted, identifying multiple novel candidates in *C. elegans* and human samples. We confirm the Pol III-dependence and noncanonical biogenesis of human miR-4521. The *mir-1829* family, miR-4521, and additional candidate miRNAs are evolutionarily young; thus, these miRNAs may represent a snapshot of miRNA loci at an early stage of their emergence.

## Results

### The *mir-1829* family is Microprocessor-independent

While investigating miRNA decay in *C. elegans* adults using a temperature-sensitive (ts) allele of the DGCR8 ortholog *pash-1* (Vieux et al. 2021; Lehrbach et al. 2012), we noticed a small portion of miRNAs that appeared to be insensitive to *pash-1* inactivation (Figure 1A, x-axis). These *pash-1*-insensitive miRNAs included both known MP-independent mirtrons (Chung et al. 2011; Jan et al. 2011; Ruby et al. 2007) and uncharacterized miRNAs such as the *mir-1829* family. To determine whether the *mir-1829* family is also MP-independent, we compared the *pash-1(ts)* data to another dataset in which RNAi and the auxin induced degron system were used together (RNAiD) to stringently deplete the MP in embryos (Dexheimer et al. 2020) (Figure 1A, y-axis). In comparing the two datasets, we observe that most miRNAs are sensitive to MP inactivation in both datasets as shown by their decreased abundance (Figure 1A). Multiple miRNAs decrease upon MP inactivation only in the adult dataset; many of these are substrates of EBAX-1-dependent miRNA decay (likely target-directed miRNA degradation) in adults but not embryos (Kotagama et al. 2024; Stubna et al. 2024) (Figure S1A); thus, their differential stability (higher in embryos) could explain their differential depletion upon MP inactivation. Other miRNAs that show differential MP sensitivity could also represent instances of differential stability or could be due to differences in the experimental systems [embryo RNAiD vs adult *pash-1(ts)*]. miRNAs that are insensitive to MP inactivation in both datasets included MP-independent mirtrons and uncharacterized miRNAs. Among the novel MP-insensitive miRNAs, the vast majority are members of the *mir-1829* family. The *mir-1829* family is comprised of four miRNA loci, *mir-1829a, mir-1829b, mir-1829c,* and *mir-4812*, with highly similar sequences, including a shared seed sequence in the 3p-derived strand. Both 5p and 3p miRNAs of these four loci are MP-independent. (A fifth locus, *F39B1.3*, bears high similarity to those of the *mir-1829* family, but does not appear to be expressed.) Thus, we hypothesized that the *mir-1829* family are representatives of a new noncanonical MP-independent class of miRNAs.

**Figure 1.**
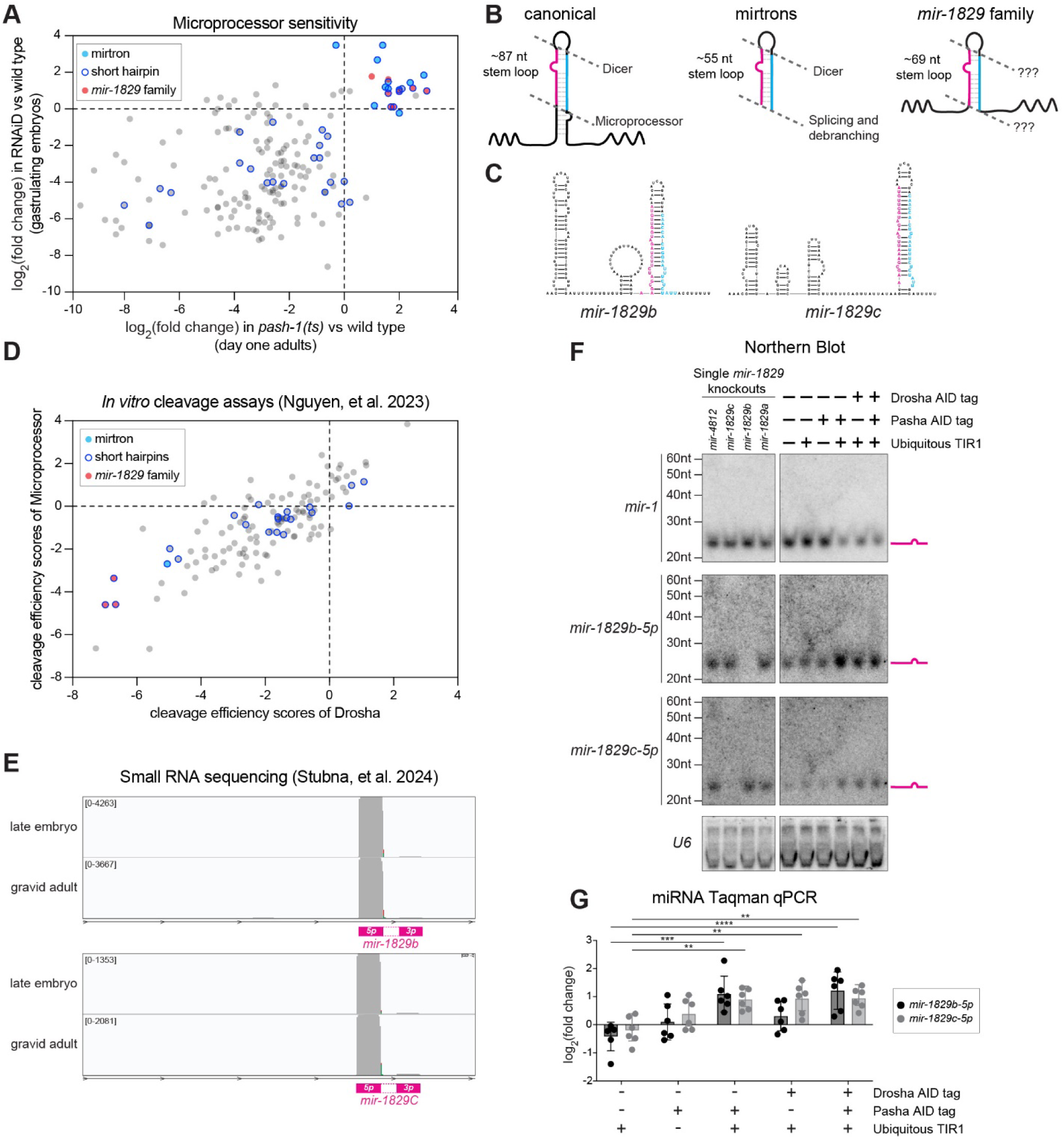
The *mir-1829* family is Microprocessor-independent. A) Comparison of log_2_(fold change) of miRNAs 24h post-upshift to restrictive temperature in *pash-1(ts)* vs. wild type (Vieux et al. 2021) (x-axis) to those in RNAiD depletion of the Microprocessor (Dexheimer et al. 2020) (y-axis). miRNAs are annotated as having a “short hairpin” if predicted structure on MirGeneDB has <32bp (Fromm et al. 2020). A small number of miRNAs are insensitive to Microprocessor depletion in both datasets, including mirtrons and the *mir-1829* family. Only miRNAs with baseMean ≥ 10 in the *pash-1(ts)* data set are plotted. B) Schematics of miRNA structures emphasizing long basal stem of canonical miRNAs and lack thereof in mirtrons and the *mir-1829* family. C) Predicted secondary structures of the *pri-mir-1829b* and *pri-mir-1829c.* D) Cleavage efficiency scores of Drosha (x-axis) or Microprocessor (y-axis) was calculated as log_2_(∑*N*_P_ + 0.1) – log_2_(∑*N*_S_ + 0.1), where *N*_P_ and *N*_S_ are normalized counts of cleaved products and pri-miRNA substrates, respectively (Nguyen et al. 2023). The *mir-1829* family is among the least favorable substrates. E) Genome browser tracks showing raw sequence reads from bias-minimized small RNA cloning from Stubna, et al. 2024. F) Northern blot of total RNA from adult animals exposed to auxin for 24h starting at the L4 stage at 20°C. *mir-1829b-5p*, *mir-1829c-5p*, and U6 probes were 5’ end-labelled, whereas *mir-1* probe was labelled according to Starfire method (Behlke et al. 2000). High stringency conditions were used for probing; see methods for details. G) Same samples used for northern blotting were assayed by RT-qPCR. miRNA expression is normalized to a small RNA control (*sn2429*), then further normalized to wild type samples. Mean and standard deviation of six biological replicates shown. Two-way ANOVA (\*\*\*\**p*<0.0001, \*\*\**p*<0.001, \*\**p*<0.01).

Canonical miRNAs are characterized by a ∼35bp/∼87nt stem loop hairpin which serves as an optimal substrate for MP cleavage (Auyeung et al. 2013; Fang and Bartel 2015; Nguyen et al. 2023; Kang et al. 2021; Kim et al. 2021; Partin et al. 2020; Jin et al. 2020) (Figure 1B). Mirtrons, which are derived from splicing, bypass the MP and serve as substrates for Dicer cleavage. Mirtrons generally have shorter hairpins than canonical miRNAs, lacking a basal stem that is characteristic of MP substrates (Westholm and Lai 2011) (Figure 1B). To determine whether newly-identified MP-independent miRNA candidates also lack a basal stem, we examined the distribution of miRNAs derived from primary transcripts containing only a “short hairpin” (<32bp) among the *pash-1(ts)* and MP RNAiD data. While “short hairpins” were present among MP-sensitive miRNAs, the MP-insensitive miRNAs were highly enriched for “short hairpins”, highlighting both known mirtrons and novel MP-independent candidates like the *mir-1829* family (Figure 1A-B).

The structure of the *mir-1829* family – along with their MP-insensitivity – suggests that they are poor MP substrates (Figure 1A-C). Consistent with this, the *mir-1829* family were among the least-efficiently cleaved pri-miRNAs in *in vitro* processing assays performed using purified *C. elegans* Drosha (Figure 1D, x-axis) or the whole MP complex (Figure 1D, y-axis) (Nguyen et al. 2023). Despite their apparently MP-independent biogenesis, the *mir-1829* family members exhibit sequencing reads from both the 5p and 3p strands and are expressed at a level comparable to canonical miRNAs (Figure 1E, Figure S1B).

We further confirmed these published sequencing datasets by examining miRNA levels via northern blotting and RT-qPCR. We used strains in which PASH-1 and/or DRSH-1 are tagged with the auxin inducible degron (AID) to deplete components of the MP for 24 h (Dexheimer et al. 2020) (Figure S1C). Using stringent northern blot conditions, we observed depletion of a canonical miRNA (*mir-1*) in samples depleted of PASH-1 and/or DRSH-1, demonstrating efficient inactivation of MP (Figure 1F). (Note that high stringency conditions were used here to distinguish between *mir-1829b-5p* and *mir-1829c-5p*. See methods for details of high stringency conditions.) In contrast, mature *mir-1829b-5p* and *mir-1829c-5p* were not diminished in samples depleted of the MP; in fact, they were modestly up-regulated (Figure 1F). (Among the *mir-1829* family, only the 5p guide strands of *mir-1829b* and *mir-1829c* were detectable by northern. We similarly focused on *mir-1829b-5p* and *mir-1829c-5p* for all RT-qPCR experiments because they were also the most robustly detectable *mir-1829* family members in these assays.) Similar to northern blots, RT-qPCR showed slightly higher levels of *mir-1829b-5p* and *mir-1829c-5p* in the same samples (Figure 1G). The increased accumulation of noncanonical miRNAs upon MP inactivation has been noted previously (Kim et al. 2016; Sheng et al. 2018; Babiarz et al. 2008) and likely reflects reduced competition from more abundant canonical miRNAs for Ago loading and/or Dicer-mediated processing. Overall, these data confirm deep sequencing data indicating that MP inactivation does not impair *mir-1829* family biogenesis. The caveat that the lack of sensitivity to MP depletion could be attributed to an unusually long half-life of the *mir-1829* family is addressed below in experiments demonstrating Dicer-dependence.

### The *mir-1829* family is germline-enriched

*C. elegans* has 19 Ago proteins, including three that are primarily dedicated to loading miRNAs: <u>A</u>rgonaute-like <u>g</u>ene 1 (ALG-1), ALG-2, and ALG-5 (Seroussi et al. 2023). ALG-1 and ALG-2 are both expressed ubiquitously in the soma, and ALG-2 is also expressed in the germline and early embryos (Aalto et al. 2018; Vasquez-Rifo et al. 2012). ALG-5 expression is restricted to the germline (Seroussi et al. 2023; Brown et al. 2017). To examine potential downstream functions of the MP-independent *mir-1829* family, we examined its Ago loading profile and tissue of expression. According to small RNA sequencing of immunoprecipitations of *C. elegans* Argonautes (IP-sRNA-seq), all members of the *mir-1829* family are highly enriched in ALG-5 IPs and modestly enriched in ALG-2 IPs (Brown et al. 2017; Seroussi et al. 2023). Knockout of ALG-1 decreases levels of multiple family *mir-1829* family members, whereas ALG-2 and ALG-5 knockout display depletion of one or two of the miRNA family strands, respectively (Seroussi, et al. 2023) (Figure S2A). Building upon these data, we sought to determine the relative distribution of the *mir-1829* family in each miRNA Argonaute via IP-RT-qPCR from strains in which ALG-1, -2, or -5 is tagged at its endogenous locus with a 3xFLAG epitope (as well as GFP or mKate2) (Aalto et al. 2018; Brown et al. 2017). We observed that *mir-1829b-5p* is evenly distributed between ALG-2 and ALG-5, whereas *mir-1829c-5p* is evenly distributed between all three miRNA Argonautes (Figure S2B).

We reasoned that our IP-RT-qPCR data could be consistent with IP-sRNA-seq data (in which higher amplitude of enrichment was observed in ALG-5 IPs than ALG-1/2 IPs) if low abundance of ALG-5 imparts higher dynamic range to ALG-5 IP enrichment scores than those for ALG-1 and ALG-2. We therefore determined the relative expression of ALG-1, ALG-2, and ALG-5 in whole animal adult samples; because each Argonaute is endogenously tagged with 3xFLAG, the anti-FLAG signal can be compared across strains to assess relative Argonaute abundance (Figure S2C, input lanes). Accordingly, ALG-1 makes up ∼60% of the miRNA Argonaute pool, whereas ALG-2 makes up ∼37% and ALG-5 a mere ∼3% in whole adult hermaphrodite samples (Figure S2D). Therefore, the dynamic range of enrichment from whole animal samples is greater for ALG-5 IPs than those of ALG-1 and ALG-2, explaining the even distribution of *mir-1829b/c* across multiple Argonautes. Although *mir-1829* family members can be detected in some somatic tissues (Alberti et al. 2018; Brosnan et al. 2021), the exclusive loading of *mir-1829b-5p* in ALG-5 and ALG-2 suggests that its expression is very strongly germline-enriched, limiting association with ALG-1 (which is not expressed in the germline). The even distribution of *mir-1829c-5p* across ALG-1/2/5 suggests that its expression is strongly enriched in the germline (given that ∼33% of the miRNA is loaded into the 3% of the Argonaute pool comprised of ALG-5), along with some expression in the soma. Additionally, germline-specific profiling of miRNAs shows that all *mir-1829* family members are highly expressed in the distal germline during the mitosis to meiosis transition (Diag et al. 2018).

Because of its germline-enriched expression, we sought to determine if the *mir-1829* family plays a role in germline function, specifically in fertility. Using CRISPR-Cas9, we generated strains that contain various combinations of *mir-1829* knockouts including single knockouts of each family member, a double mutant of *mir-1829b* and *mir-1829c*, a triple mutant of *mir-1829b/c* and *mir-4812*, and a quadruple knockout of all four family members (Figure S2E). Reduced brood size and embryonic lethality were initially observed in the single *mir- 1829a* deletion [*mir-1829a(del)*] and the full quadruple knockout of all members of the *mir-1829* family [*mir-1829 family(del)*] at elevated temperatures (Figure S2E). However, these phenotypes were not rescued by a second round of CRISPR that restored the *mir-1829a* sequence [*mir- 1829a(rescue)*] and thus are due to off-target effects of CRISPR (Figure S2E). The *mir-1829* family members reside in introns of protein coding genes (see also below), and the host gene of *mir-1829a*, *gas-1*, is required for normal fertility (Morgan and Sedensky 1994). We therefore redesigned the *mir-1829a* deletion to minimize effects on *gas-1* expression by deleting the whole *mir-1829a* host intron and fusing the flanking exons together [*mir-1829a(intron del)*]; this strain did not show abnormal brood or embryonic viability phenotypes (Figure S2E). Finally, we used *mir-1829a(intron del)* to generate another quadruple mutant strain in which all four family members are deleted [*mir-1829 family(null)*], and this was also superficially wild type. Therefore, the *mir-1829* family is dispensable for normal fitness in laboratory conditions (Figure S2E).

ALG-5 has a demonstrated role in promoting the mitosis to meiosis transition in the *C. elegans* germline, evidenced by strong enhancement of the *glp-1(gf)* tumorous phenotype by *alg- 5(null)* (Brenner et al. 2022) (Figure S2F). Because of the expression of the *mir-1829* family in the distal germline and its loading in ALG-5, we examined its role in this context. We observed that the quadruple knockout of the *mir-1829* family does not modify the *glp-1(gf)* phenotype (Figure S2F); thus, it is not the primary miRNA guiding ALG-5’s function in this context. Overall, the biological function of the *mir-1829* family remains to be determined.

### ALG-5 represses expression of a tethered mRNA

Because of its shared miRNA binding repertoire with ALG-1/2, ALG-5 likely acts as a repressor of its targets, though it has also been proposed to activate targets (Liontis et al. 2023). Since ALG-5’s targets remain to be determined, we tested the functionality of ALG-5 using a tethering assay. Briefly, a single-copy GFP-histone transgene containing three boxB RNA hairpins in its 3 UTR is expressed throughout the germline (Aoki et al. 2021). In this background, we fused a 3xFLAG tag and λN22 peptide to the N-terminus of ALG-5 by CRISPR (or 3xFLAG alone as a negative control background). Because the reporter is prone to silencing, we simultaneously inactivated *mut-2* by CRISPR in both strains. Due to the strong interaction of the λN22 peptide with boxB hairpins, 3xFLAG-λN22::ALG-5 but not 3xFLAG::ALG-5 should be tethered to the reporter transcript allowing us to assess the impact of ALG-5 association with an mRNA by comparing the two strains. Imaging of these two strains shows lower expression of the GFP-histone reporter in 3xFLAG-λN22::ALG-5 than 3xFLAG::ALG-5 in the distal germline of L4 and adult animals (Figure 2A-C). Thus, this *in vivo* tethering assay suggests that ALG-5 acts similarly to other miRNA Agos by repressing associated transcripts.

**Figure 2.**
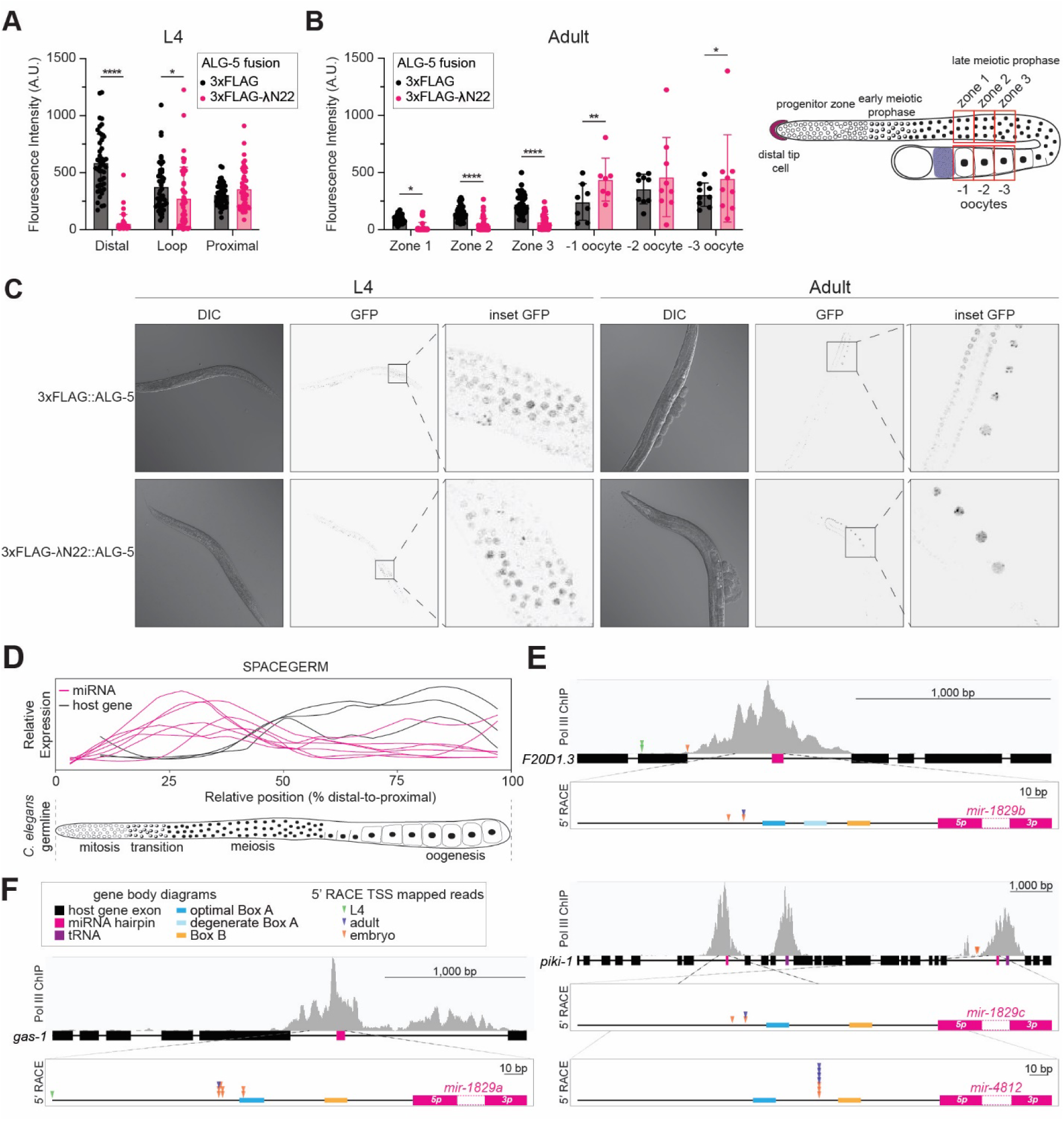
ALG-5-mediated reporter repression and *mir-1829* family expression both peak in the distal germline. A) Measured GFP fluorescence intensity from five nuclei per region (distal, loop, and proximal) across nine L4-staged animals, grown at 20°C expressing a germline single-copy GFP::histone transgene with three boxB RNA hairpins in its 3 UTR. CRISPR- generated animals expressing ALG-5 fusion proteins were tagged with either 3xFLAG and λN22 (experimental) or 3xFLAG alone (control) in a *mut-2(null)* background. B) Measured relative GFP fluorescence intensity in Day 1 adult worms (same strains as in panel A). Average fluorescence intensity was measured from six regions of interest (shown in right schematic). Zones 1–3 correspond to regions of the distal gonad across from the –1, –2, and –3 oocytes, respectively. Five nuclei were measured in each zone in the distal germline per nine adult worms. A-B) Mean and SD shown. Two-way ANOVA (****p<0.0001, **p<0.01, *p<0.05). C) Representative images of A and B. D) Schematic of relative expression levels of the *mir-1829* family and its host genes in the *C. elegans* germline. Data from SPACEGERM (Diag et al. 2018) indicates that the *mir-1829* family (pink) peaks in expression in the distal germline, whereas its host genes (black) peak in expression in the proximal germline. E-F) Prominent Pol III ChIP-seq peaks (Araya et al. 2014) reside at the *mir-1829* loci (pink). Arrowheads: 5’ RACE reads cloned from RNA of indicated stage. Putative Pol III promoter sequence motifs upstream of the miRNA precursors are indicated.

### *The mir-1829* family is differentially transcribed from its host genes

The four *mir-1829* family loci reside in the introns of three protein coding genes of apparently unrelated function (*gas-1*, *piki-1*, and F20D1.3). As mentioned above, data from the SPACEGERM (<u>Spa</u>tial *<u>C</u>aenorhabditis **e**legans* <u>g</u>ermline **e**xpression of m<u>R</u>NA & <u>m</u>iRNA) atlas indicate that the *mir-1829* family peak in expression in the distal germline (Diag et al. 2018) (Figure 2D). Curiously, the host genes peak in expression in the proximal gonad, suggesting differential expression of the miRNAs and their host genes (Figure 2D). Published ChIP-seq data of Pol III subunit RPC-1 further supports the idea that the *mir-1829* family is transcribed independently from its host genes and by Pol III (Araya et al. 2014); the *mir-1829* loci (*mir-1829* precursors shown in pink) are marked by prominent Pol III ChIP peaks, and these peaks are just as prominent as those at tRNA loci (purple boxes) (Figure 2E-F).

To investigate whether the differential expression between host gene and miRNA is due to differential transcription, we performed 5’ Rapid Amplification of cDNA Ends (RACE) on RNA isolated from embryo, L4, and adult tissues using primers that bind in the apical loop of the miRNA hairpins. For each of the *mir-1829* family loci, multiple 5’ RACE reads mapped within the host intron of the miRNA, suggesting alternative proximal transcription start sites of transcripts independent of the protein-coding host genes (Figure 2E-F, Table S3). For three of the four members of the *mir-1829* family (*mir-1829a-c*), multiple 5’ RACE reads mapped consistently 101-103bp upstream of the precursor, which further suggests the *mir-1829* family members share conserved regulatory elements that drive their independent transcription.

### *The mir-1829* family is transcribed by Pol III

Because results of 5’ RACE together with published Pol III ChIP suggest that the *mir- 1829* family may arise from independent Pol III transcriptional units, we sought additional evidence of their Pol III dependence. Type II Pol III transcripts (e.g. tRNAs) are defined by having Box A and Box B motif promoter elements located downstream of the transcription start sites (TSS) (Dieci et al. 2007) (Figure 3A). For each *mir-1829* family member, we searched upstream of the precursor for the Box A and Box B consensus sequence motifs previously defined for *C. elegans* (Ikegami and Lieb 2013; Stutzman et al. 2020). We determined that all four members have a putative Box A and Box B motif ∼79-87bp and ∼35-42bp upstream of the precursor, respectively. The positions of the motifs are downstream of the prominent -102bp TSS identified by 5’ RACE for *mir-1829a*, *mir-1829b*, and *mir-1829c* (Figure 3A). Additionally, all the *mir-1829* family loci have 5Ts immediately downstream of the precursor (3-4bp downstream of the 3p arm), and another stretch of 4-5Ts further downstream. These features further support our hypothesis that these are Pol III transcripts since typical Type II Pol III genes have terminator sequences of 5Ts immediately downstream of the gene body (Ikegami and Lieb 2013).

**Figure 3.**
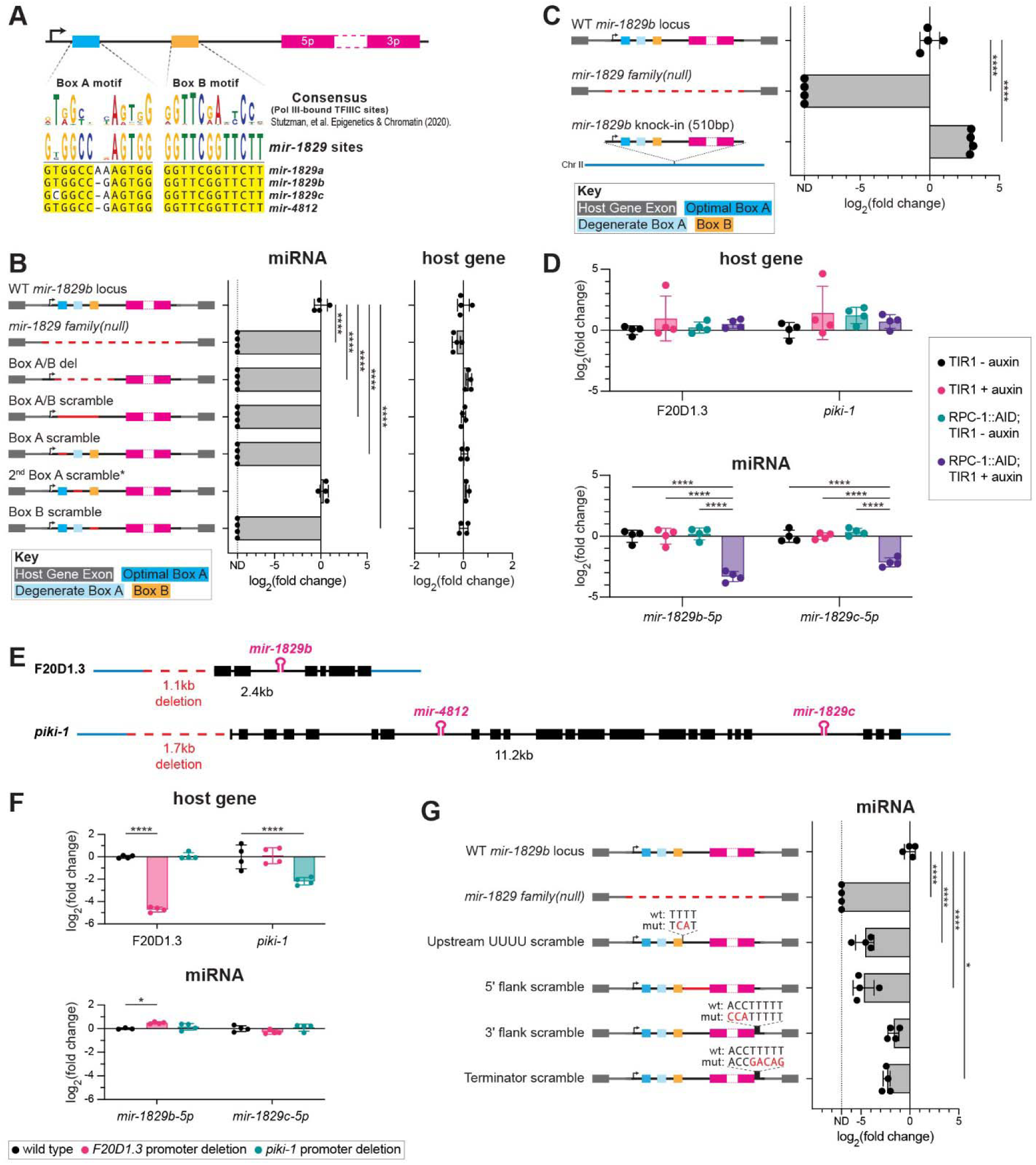
*mir-1829* family members are transcribed by Pol III. A) MUSCLE alignment of the *mir-1829* family putative Pol III promoter sequence motifs. B) Schematic of CRISPR alleles generated at the *mir-1829b* locus (left) and qPCR of resulting miRNA and host gene expression (right) from young adult samples in wildtype background, grown at 25°C. C) Schematic of genomic sources of *mir-1829b* (left), including re-integration of a 510bp minimal *mir-1829b* transcriptional unit in a *mir-1829 family(null)* mutant background (bottom left). qPCR of *mir- 1829b* expression from young adult samples grown at 25°C (right). D) qPCR of miRNAs and host gene transcripts in AID-tagged RPC-1 strain with ubiquitous TIR1 (MCJ666) or TIR1 alone (MLC1040). Auxin treatment was at 20°C for 24h, beginning at the L4 stage. E) Schematics of promoter deletions for two host genes of the *mir-1829* family. F) qPCR in host gene promoter deletion samples from young adult samples grown at 25°C. G) Schematic of CRISPR-generated *mir-1829b* alleles reintroduced into *mir-1829 family(null)* (left) and qPCR of resulting miRNA (right) from young adults, grown at 25°C. B-D, F-G) miRNA expression is normalized to a small RNA control (*sn2429*) and then further normalized to wild type levels. Host gene expression was normalized to GAPDH (*gpd-1*) and then to wild type levels. Mean and SD shown. Two-way ANOVA. \*\*\*\**p*<0.0001, \**p*<0.05.

To determine whether the Box A/B motifs are required for *mir-1829* biogenesis, we performed promoter bashing experiments for *mir-1829b*. As a control, we used the *mir-1829 family(null*) mutant in which all the family members are deleted; in this background, a 312bp deletion that removes *mir-1829b* abolishes expression of the miRNA without disrupting the host gene transcript, F20D1.3 (Figure 3B). The *mir-1829b* locus contains a putative Box A motif, a second degenerate Box A motif, and a putative Box B motif. We first deleted all the promoter motifs, an 82bp deletion, keeping the TSS intact (Box A/B del). This mutant resulted in abolished expression of the miRNA without disrupting the host gene transcript. To ensure that any changes in expression observed were not due to disruption of the secondary structure, we designed a new mutant (Box A/B scramble) in which we scrambled the sequence within a 62bp window, removing the Box A and Box B sequence motifs while maintaining secondary structure of the transcript as predicted by RNAfold (Lorenz et al. 2011; Gruber et al. 2008). Similar to the Box A/B del mutant, we observed abrogated miRNA expression, without disruption of the host gene transcript (Figure 3B).

We next mutated each motif individually. To maintain structure for the Box A scramble mutant, we introduced additional mutations into the degenerate Box A motif, since they form a hairpin according to the RNAfold prediction. In this Box A scramble mutant, we observed abrogated *mir-1829b-5p* expression (Figure 3B). We separately mutated only the degenerate Box A motif in a manner that is not predicted to maintain secondary structure (2^nd^ Box A scramble); this resulted in no change in expression of *mir-1829b-5p* (Figure 3B). Lastly, in the Box B scramble mutant, we maintained predicted structure while disrupting the sequence of the Box B motif and observed loss of *mir-1829b-5p* expression. Together, these results indicate that the Box A and Box B motifs, individually, are each required for *mir-1829b* expression, whereas the degenerate Box A motif is not.

Building on the promoter bashing experiments, we attempted to express the predicted *mir-1829b* locus from an ectopic location. We re-integrated a 510bp minimal transcriptional unit – spanning 125bp upstream of the TSS to 222bp downstream of the precursor – into the *mir- 1829 family(null)* background at a known safe harbor locus (the position of MosSCI locus *ttTi5605* in Chromosome II). Not only did we observe expression of *mir-1829b-5p* in this rescue strain, but expression was increased nearly eight-fold compared to wild type (Figure 3C). Northern blot confirmed that the knock-in locus generated a distinct *mir-1829b-5p* small RNA species (Figure S3); since germline-specificity of this species has not been tested, broader tissue expression or reduced interference from host gene transcription could underlie the higher expression levels (see also Discussion). Overall, the observed expression of the minimal transcriptional unit demonstrates that the 510bp fragment contains all the required regulatory elements necessary for *mir-1829b* expression.

To further test the model that the *mir-1829* family is a Pol III transcript, we depleted Pol III using the AID system. To do this, we tagged the Pol III catalytic subunit RPC-1 with a 3xFLAG::AID tag using CRISPR in a strain that expresses TIR1 ubiquitously, resulting in efficient RPC-1 depletion upon auxin treatment (Figure S4). We assessed expression of *mir- 1829b-5p*, *mir-1829c-5p*, and their respective host gene transcripts after 24 h of auxin treatment using RT-qPCR. Depletion of Pol III by auxin treatment of the AID-tagged RPC-1 strain did not impact levels of the Pol II-transcribed host gene transcripts but strongly abrogated expression of *mir-1829* family members (Figure 3D). In contrast, *mir-1829b/c* levels were unaffected in RPC-1 degron strains lacking auxin treatment, signifying that tagging RPC-1 alone does not disrupt its function (Figure 3D). Furthermore, the significant depletion of *mir-1829b/c* was not due to off target effects of auxin treatment or TIR1 expression, as observed by the lack of miRNA depletion in strains that do not contain tagged RPC-1 (Figure 3D – “TIR1 +/- auxin”). Overall, depletion of RPC-1 along with the mutation of BoxA/B motifs support the model that the *mir- 1829* family is transcribed by Pol III.

To determine if Pol II also plays a role in the transcription of the *mir-1829* family, we deleted the promoter regions of the host genes with the rationale that this would remove Pol II occupancy from these loci. We deleted 1.1kb and 1.7kb upstream of the start codon for F20D1.3 and *piki-1*, respectively (Figure 3E). We observed strong reduction of host gene expression levels in their respective mutants via RT-qPCR, suggesting that we effectively removed Pol II occupancy (Figure 3F). In the *piki-1* promoter deletion mutant, there was no change in *mir- 1829c-5p* levels. In the F20D1.3 promoter deletion mutant, expression of *mir-1829b-5p* was slightly increased (Figure 3F). We hypothesize that there may be slight competition between Pol II and Pol III for the F20D1.3*/mir-1829b* locus, since deletion of the F20D1.3 promoter region (Figure 3F) or ectopically expressing *mir-1829b* from another locus (Figure 3C) results in increased expression of *mir-1829b-5p*. Together, these results validate the model that the *mir- 1829* family is primarily transcribed by Pol III, with no detectable contribution from Pol II-driven host gene transcription.

Having demonstrated the dependence of *mir-1829b* on Pol III and its promoter elements, we returned to the *mir-1829b* locus to assess the role of additional *cis* elements. For these experiments, variants of the *mir-1829b* gene were knocked back in to its genomic locus in the context of the *mir-1829 family(null)* background; this approach facilitated genotyping for isolation of very small sequence changes. All mutations were designed to preserve predicted secondary structure of pri-*mir-1829b*. First, we tested whether 4Ts present upstream of the precursor suppress productive *mir-1829b* transcription by causing early termination. This is not the case since mutating this run of Ts decreased *mir-1829b* expression (Figure 3G, Upstream UUUU scramble). Next, we tested whether the regions flanking the precursor in the primary transcript are important for *mir-1829b* biogenesis. This was true of the 33nt 5 -flanking sequence whose mutation reduced *mir-1829b* level, but not of the 3nt 3 flank whose mutation had an insignificant effect (Figure 3G, 5’ and 3’ flank scramble). Finally, we tested the importance of the first run of Ts after the precursor; mutating these five residues reduced *mir- 1829b* expression, but did not abolish it (Figure 3G, Terminator scramble). This suggests that these 5Ts play a role in biogenesis – likely by promoting Pol III termination – and the intact downstream 4T tract may support the residual *mir-1829b* expression. The decreased expression in this mutant may suggest that a very short flank 3 of the precursor promotes biogenesis.

### *mir-1829* family biogenesis requires Dicer, but not DRH-1, RDE-4 or NSUN-2

Having demonstrated that the *mir-1829* family is Pol III-transcribed and MP-independent, we sought to further elucidate its biogenesis. To do so, we employed the AID system to determine the dependence of *mir-1829* biogenesis on Dicer (DCR-1). DCR-1 was tagged with 3xFLAG-AID at its endogenous locus and crossed into a background expressing TIR1 ubiquitously or only in the germline, resulting in dramatic or moderate depletion of overall DCR-1, as expected (Figure S4). We first performed northern blotting to examine the abundance of the mature miRNA species and the accumulation of the precursor. As a control, we probed for the canonical miRNA *mir-1*, which is primarily expressed in the pharynx and muscle (Brosnan et al. 2021; Simon et al. 2008; Xu et al. 2024; Gutiérrez-Pérez et al. 2021). We observed reduction of mature *mir-1* and accumulation of its precursor in the samples where Dicer is depleted ubiquitously but not in the case of germline-only Dicer depletion (Figure 4A). To achieve maximal signal for *mir-1829b/c,* low stringency conditions were used that do not distinguish between *mir-1829b-5p* and *mir-1829c-5p* when using a *mir-1829b-5p* probe, as demonstrated by detection of pure synthetic miRNA species corresponding to both sequences (Figure 4A, right side, see methods for low stringency conditions). We observe decreased *mir-1829b/c* expression in all the Dicer-depleted samples, including those in which Dicer is depleted only in the germline, consistent with strong germline enrichment of the *mir-1829* family (Figure 4A). Despite low stringency probing conditions, the *mir-1829* family is still near the limit of detection (Figure 4A). This low signal may contribute to our failure to detect accumulation of the precursor species of *mir-1829b/c* as expected upon Dicer depletion.

**Figure 4.**
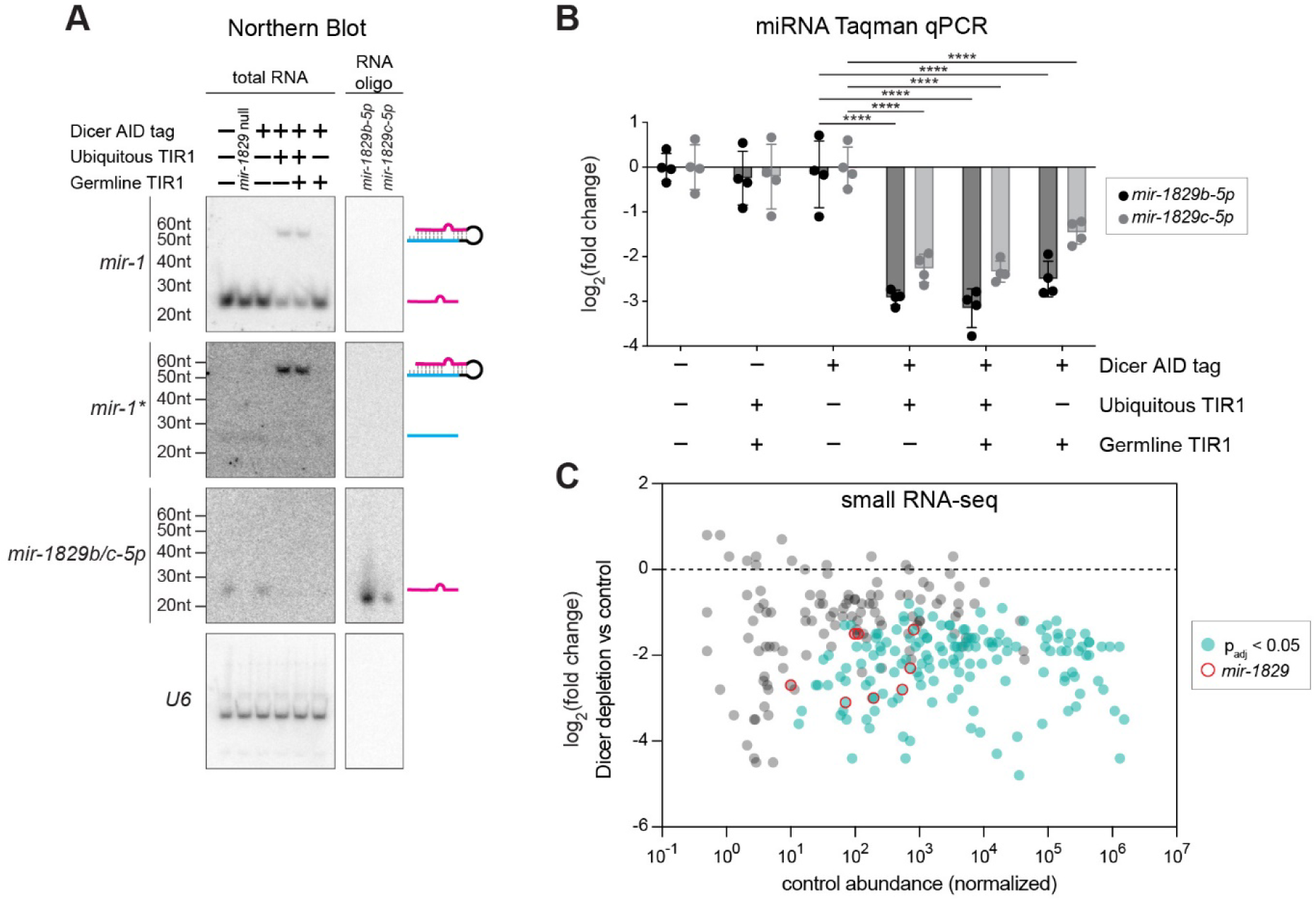
The *mir-1829* family is Dicer-dependent. (A-C) All samples were exposed to auxin for 24h starting at the L4 stage, then harvested as adults at 20°C. A) Northern blot using low stringency conditions (see methods for details of low stringency conditions). Right panels are lanes loaded with synthetic RNA oligos, demonstrating recognition of both *mir-1829b-5p* and *mir-1829c-5p* with the *mir-1829b-5p* probe under these conditions. B) miRNA qPCR normalized to a small RNA control (*sn2429*) and then further normalized to levels in control that only carries Dicer AID tag (UY212). Mean and SD shown. Two-way ANOVA. \*\*\*\**p*<0.0001 C) Small RNA-seq. MA plot showing abundance in control (DCR-1::AID tag alone, UY212) on x-axis and log_2_(fold change) in DCR-1::AID; ubiquitous TIR1 (MCJ387) compared to UY212 on y-axis. miRNAs showing significant sensitivity to Dicer depletion are shown in blue-green, and the *mir-1829* family is highlighted in red open circles. Three biological replicates of each genotype were analyzed using DESeq2.

Due to the limited detection of the *mir-1829* family by northern blotting, we further assessed miRNA levels using RT-qPCR. We observed significant reduction of both *mir-1829b- 5p* and *mir-1829c-5p* in samples in which Dicer is ubiquitously depleted (Figure 4B). Furthermore, the level of reduction in *mir-1829b-5p* in germline-only Dicer depletion samples nearly mirrors the level seen in the ubiquitously depleted samples (Figure 4B), further supporting germline-restricted expression of *mir-1829b-5p*. We observe milder depletion of *mir-1829c-5p* in the germline Dicer depletion samples compared to depletion in the whole worm (Figure 4B). This result further supports that *mir-1829c* is expressed in the both the germline and the soma as noted above (Figure S2B).

To further assess Dicer dependence, we performed sRNA-seq of samples in which Dicer is depleted ubiquitously or in the germline (Tables S6-S7). Overall, almost all miRNAs appear to be Dicer-dependent – many of which are statistically significant after spike-in normalization (*p*_adj_ < 0.05) (Figure 4C, Table S6). The *mir-1829* family behaves similarly to canonical miRNAs in that it is sensitive to Dicer depletion (Figure 4C, red circles). Consistent with all results discussed above, we also observed depletion of the *mir-1829* family members when Dicer was depleted in the germline alone, along with depletion of a small set of germline-enriched canonical miRNAs (Figure S5, Table S7).

To process long dsRNA substrates, *C. elegans* Dicer works in concert with two cofactors, double stranded RNA (dsRNA) binding protein RDE-4 and RIG-I-like receptor DRH-1 (Tabara et al. 2002; Duchaine et al. 2006; Lu et al. 2009; Consalvo et al. 2024). Since the *mir-1829* family is noncanonically processed but requires Dicer, we tested the role of RDE-4 and DRH-1 in *mir-1829* biogenesis using knockout strains. From RT-qPCR experiments, we observed no change in *mir-1829b/c* in the *rde-4* and *drh-1* mutant strains compared to wild type (Figure S6A). Thus, Dicer functions in *mir-1829* maturation independent of RDE-4 and DRH-1.

We also investigated the role of *nsun-2* and multiple cellular nucleases in *mir-1829* biogenesis. Vault RNAs can be further processed by Dicer to produced small RNA fragments (svRNAs) that function in RNAi when loaded into Ago (Persson et al. 2009). In cell lines, NSun2, an RNA methyltransferase that modifies cytosine, methylates vault RNA to prime them for dicing (Hussain et al. 2013). To determine if the *C. elegans* homolog *nsun-2* plays a role in *mir-1829* biogenesis, we introduced a 3xFLAG-AID tag to the C-terminus of NSUN-2 by CRISPR. After 24 h auxin treatment in a ubiquitous TIR1 background, although NSUN-2 levels decreased approximately 90% (Figure S4), *mir-1829b/c* levels remained unchanged (Figure S6B). Thus, *nsun-2* does not appear to play a role in *mir-1829* family biogenesis. To determine the role of exonucleases in *mir-1829* family biogenesis, we reanalyzed data from a study in which most core cellular nucleases were knocked down by RNAi, followed by small RNA sequencing (sRNA-seq) (Vieux et al. 2021). None of these RNAi conditions altered *mir-1829* family member abundance, suggesting that these exonucleases are also dispensable for *mir-1829* family biogenesis (Figure S6C).

### The 3**’** end of the *mir-1829* family precursor is likely generated in the nucleus

Although they differ in biogenesis of their precursor, mirtrons are similar to the *mir-1829* class of miRNAs in their MP independence and Dicer dependence. Mirtron biogenesis is counteracted by untemplated 3’ end uridylation of the precursor species, which destabilizes these intermediates (Bortolamiol-Becet et al. 2015; Reimão-Pinto et al. 2015). Activity of terminal nucleotidyl transferases on miRNA precursor substrates can be detected through sequencing of mature miRNA populations since the untemplated nucleotide additions (tails) can persist after Dicer-mediated maturation. In particular, tailing that is strongly skewed toward 3p-derived miRNA strands is likely a signature of an enzyme that acts on miRNA precursor substrates; such activity would not result in tailing of 5p-derived miRNA strands whose 3’ ends are not available for modification until after Dicer-mediated precursor cleavage. We previously observed that miRNA A-tailing in *C. elegans* is strongly skewed toward 3p-derived miRNAs, suggesting that an adenyltransferase acts on miRNA precursors (Vieux et al. 2021). To determine whether *mir- 1829* family biogenesis is regulated by tailing of the precursor, similar to mirtrons, we examined the sequences of the mature strands for evidence of tailing (which may have been deposited on either precursor or mature miRNA species). We observed various tailed species of *mir-1829* family members with a clear enrichment of adenylation on the 3p-derived strands, suggestive of adenylation of the miRNA precursors (Figure 5A). Sequencing reads of *mir-1829b/c-3p* (which share an identical reference sequence and displayed the highest level of adenylation) show 3’ trimmed isoforms and reveal that adenylation occurs on an isoform that is 1nt shorter than the reference (Figure 5B). This adenylation is not predicted to change the structure of the miRNA precursor from the reference secondary structure; either an unmodified or a precursor in which the terminal base is substituted with A are predicted to have 1nt unpaired a the 5 end and 3nt unpaired at the 3 end (see Figure 1C). Although the trimmed 3p isoforms suggest that an exonuclease modifies or matures the 3 end of the precursor, no exonuclease is yet identified to play a role in biogenesis (Figure S6C).

**Figure 5.**
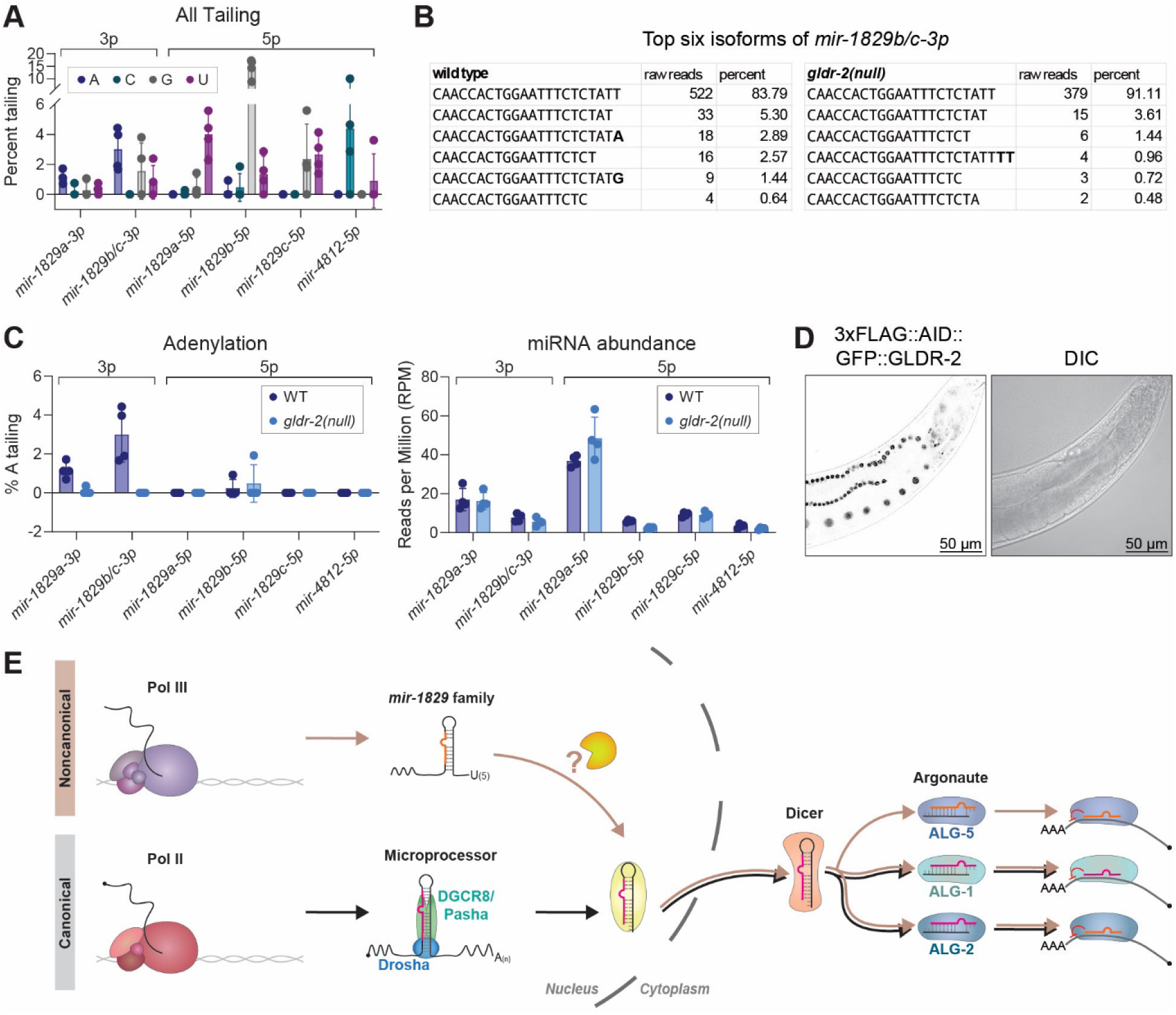
The *mir-1829* family primary transcripts are likely cleaved in the nucleus. A) Percent untemplated 3′ nucleotide additions (tailing) of the *mir-1829* family members in wild type. B) Raw read counts across four biological replicates showing top six isoforms of *mir- 1829b/c-3p.* C) Percent A-tailing in wild type and *gldr-2(null)* (top). Abundance of each mature species shown below (bottom). (A, C) All *mir-1829* family member strands with >1RPM average abundance are shown. Raw data reanalyzed from Vieux, et al. 2021. D) Representative images of 3xFLAG::AID::GFP::GLDR-2. E) Model for new miRNA biogenesis pathway of the *mir-1829* family in contrast to canonical miRNA biogenesis.

The 5p-derived strands also displayed tailed isoforms (which must arise from modification of mature miRNAs), including uridylation of *mir-1829a-5p* and *mir-1829c-5p*, cytidylation of *mir-4812-5p*, and guanylation of *mir-1829b-5p* (Figure 5A). Terminal guanylation and cytidylation of miRNAs are rarely observed in animals; uridylation is the most frequently observed modification of *C. elegans* mature miRNAs (Vieux et al. 2021). A functional role for tailing of mature miRNAs has not yet been established in *C. elegans*, and because the modifications of 5p strands differ across the family members, we did not explore a potential role in biogenesis.

Since GLDR-2 is responsible for A-tailing of miRNA precursors in *C. elegans*, we reanalyzed published data to examine if GLDR-2 is also responsible for tailing of the *mir-1829* family (Vieux et al. 2021). We observed loss of A-tailing on 3p arms of *mir-1829* family members in *gldr-2(null)* mutants (Figure 5C) (Vieux et al. 2021). To determine if A-tailing is involved in *mir-1829* biogenesis, we evaluated mature miRNA levels in the presence and absence of GLDR-2. We observed no change in the levels of expression, suggesting that, although *mir-1829* precursors can be adenylated by GLDR-2, A-tailing is dispensable for *mir- 1829* biogenesis (Figure 5C) (Vieux et al. 2021).

The A-tailing of the 3p arms of the *mir-1829* family suggests that a precursor-like intermediate (in which the 3’ end of the 3p arm is available for tailing) precedes Dicer-mediated maturation of the mature duplex, despite our inability to detect this intermediate on northern blots. We reasoned that localization of GLDR-2 may further inform the order of events of *mir- 1829* biogenesis. A GFP tag fused to GLDR-2 at its endogenous locus reveals that GLDR-2 specifically localizes to the nucleus (Figure 5D). This suggests that the cleavage event(s) that gives rise to the miRNA precursor – or at least maturation of the precursor 3’ end – occurs in the nucleus prior to Dicer-mediated cleavage in the cytoplasm (Figure 5E). Thus, we have delineated a new miRNA biogenesis pathway, as exemplified by *mir-1829* family, that involves the transcription of an atypical primary miRNA transcript by Pol III, initial MP-independent maturation steps in the nucleus, followed by Dicer-mediated cleavage in the cytoplasm (Figure 5E).

### Noncanonical miRNA biogenesis extends beyond the *mir-1829* family

Having defined atypical features of the *mir-1829* family’s biogenesis, we sought to extend these findings by identifying additional noncanonical miRNAs. First, we re-examined datasets in which MP is inactivated (Vieux et al. 2021; Dexheimer et al. 2020). For these analyses, we used uncurated miRBase annotations (Kozomara et al. 2019), rather than MirGeneDB which stringently filters for features of canonical miRNAs and evolutionary conservation (Fromm et al. 2020). Among MP-insensitive miRNAs, we observed many annotated mirtrons as expected (Figure 6A, Tables S10-11) (Chung et al. 2011; Jan et al. 2011; Ruby et al. 2007). In both MP inactivation datasets, we also observed that *mir-8196a-3p, mir- 8196b-3p*, and *mir-8199-5p* were not depleted (Figure 6A, Tables S10-11) (Vieux et al. 2021; Dexheimer et al. 2020). In contrast, all of these were decreased upon Dicer depletion over the same time scale (24 h), demonstrating that a long half-life for these small RNAs is not responsible for their perdurance in the setting of MP inactivation (Figure 6C, Table S10). Based on its sequence and loading into ERGO-1 (Seroussi et al. 2023), *mir-8199-5p* appears to be a misannotated 26G class endo-siRNA. *mir-8196a/b-3p* are more intriguing because they are bound in part by ALG-1 and ALG-2 (Seroussi et al. 2023), and therefore may function as noncanonical miRNAs. *mir-8196b* lies 65bp downstream of a recently annotated 21U piRNA on the X chromosome (Seroussi et al. 2023) (Figure S5A). Similarly, *mir-8196a* similarly lies 71bp downstream of an unannotated 21U piRNA, also on the X chromosome; the upstream RNA meets the requirements of a 21U based on its sequence, loading in PRG-1, and depletion in *prg- 1^-/-^* (Figure 6B). This consistent proximity to a 21U locus raises the possibility that this relative position may contribute to *mir-8196a/b* transcription or biogenesis. Consistent with this, *mir- 8196a/b* are germline-enriched, as evidenced by their significant reduction in samples in which Dicer is depleted only in the germline (Table S12).

**Figure 6.**
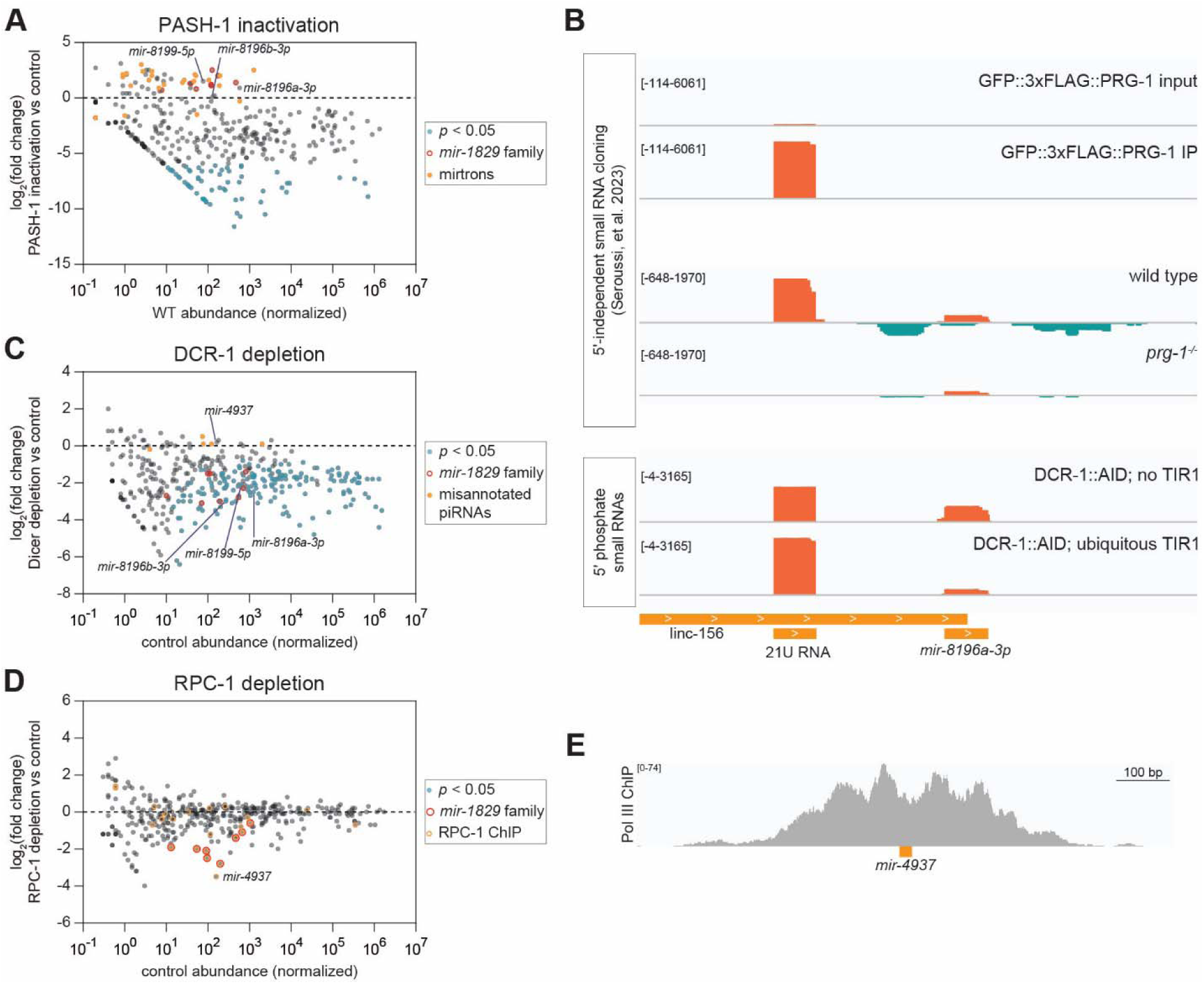
Noncanonical miRNA biogenesis pathway extends beyond the *mir-1829* family. A, C, D) Small RNA-seq results in samples depleted for 24h of functional (A) PASH-1 (C) Dicer (DCR-1) or (D) RPC-1. MA plots show log_2_(fold change) in experimental vs control on the y-axis, with normalized control abundance on the x-axis. [Controls are A) wild type (N2), C) auxin-treated DCR-1::AID tag alone (UY212), and D) auxin-treated TIR1 (MLC1040).] All miRBase annotations are plotted, and relevant noncanonical small RNAs are highlighted. B) Genome browser tracks showing proximity of *mir-8196a* to an upstream unannotated 21U RNA. 5′-independent small RNA cloning following polyphosphatase treatment also shows 22G RNAs generated from the locus. *mir-8196a* but not the 21U shows Dicer-dependence. E) Genome browser track showing RPC-1 (a Pol III subunit) ChIP at the *mir-4937* locus.

Multiple Dicer-independent small RNAs were also detected (Figure 6C, Tables S8,10-11). These include misannotated 21U piRNAs that persist in mirBase that were recently noted (Seroussi et al. 2023). Thus, these small RNAs act as positive controls for Dicer-independence, similarly to mirtrons for MP independence. We also find that *mir-5549-3p* is an additional misannotated 21U RNA encoded on chromosome III; it is 21-nt in length with U in the 5’ position, bound by PRG-1, and depleted in *prg-1^-/-^* (Seroussi et al. 2023) (Figure S5B). Finally, we identify *mir-4937* as a Dicer-independent small RNA (see more below).

We also sought to further define Pol III-dependent miRNAs. To this end, we performed sRNA-seq on samples in which RPC-1 was depleted using the AID system and overlapped these data with published Pol III ChIP-seq data to identify bona fide Pol III-transcribed miRNAs (Figure 6D, Tables S9-11). The strongest candidate is again *mir-4937*, which is RPC-1- dependent and has prominent Pol III ChIP peaks (Figure 6E). Consistent with these regulatory elements and Pol III-dependence, a previous study noted homology between *mir-4937* and a tRNA locus, mostly within putative Box A and Box B motifs (Roberts et al. 2013). Although *mir-4937* did not meet our stringent criteria for defining novel MP-independent small RNAs, its depletion in *pash-1(ts)* is marginal and not called as significant (Table S10), and its predicted secondary structure suggests that its primary transcript would make a poor MP substrate (Figure S7C). Thus, *mir-4937* is likely a Pol III-transcribed MP- and Dicer-independent small RNA. *mir- 4937* was not robustly immunoprecipitated with any of the *C. elegans* Argonautes by Seroussi, *et al*. (all average IP <0.2 reads per million) (Seroussi et al. 2023); thus, its functional class remains unclear. Nonetheless, previous studies identified it as a small RNA that increases during *C. elegans* aging (De Lencastre et al. 2010).

### Human miR-4521 biogenesis is noncanonical and Pol III-dependent

To determine whether miRNAs can derive from independent Pol III transcripts outside of *C. elegans,* we also examined ChIP datasets of Pol III subunits in three human cell lines: Hela, K562, and HEK293 (Oler et al., 2010; Dunham et al., 2012; Gerber et al., 2020, respectively). Manual curation of HEK293 RPC62 ChIP confirmed that miR-4521 and miR-4638 had strong Pol III ChIP signals (Figure 7A). Whereas miR-4638 is too lowly expressed to assess its biogenesis requirements, miR-4521 shows evidence of Drosha and Dicer independence in HEK293 cells (Figure S7D) (Rybak-Wolf et al. 2014).

**Figure 7.**
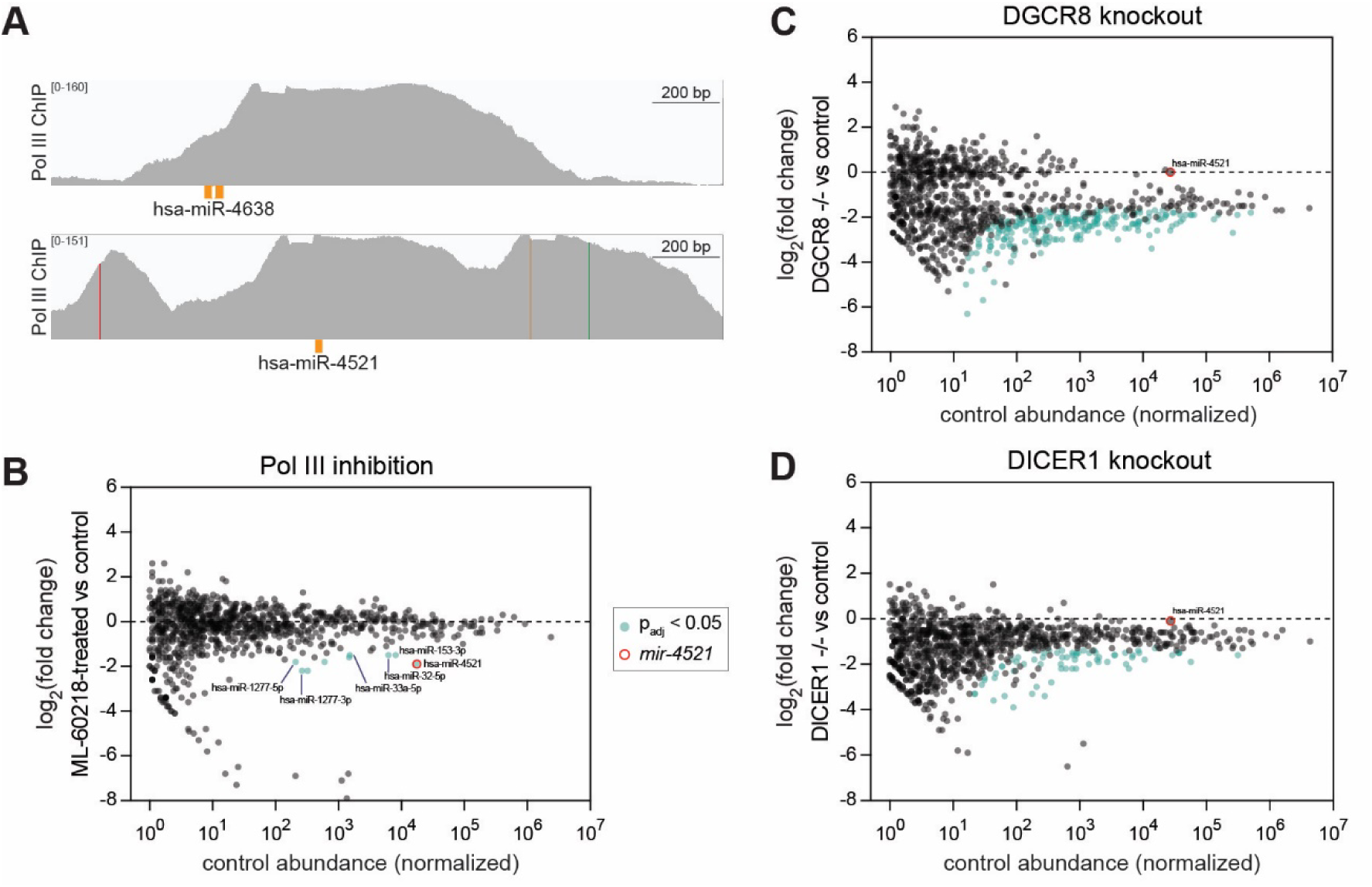
Human small RNA miR-4521 is Pol III-dependent and DGCR8- and Dicer-independent. A) Genome browser tracks showing RPC62 (Pol III) ChIP in HEK293 cells at the *mir-4638* and *mir-4521* loci. B) Small RNA-seq results from iCas9 RKO cells treated with Pol III inhibitor ML-60218 for 24h compared to vehicle-treated (DMSO) control. C-D) Small RNA-seq results from cells grown for four days after induction of guide RNAs targeting (C) DGCR8 or (D) Dicer, compared to control iCas9 RKO cells. miRNA reads are normalized to spike-ins. B-D) miRNAs with baseMean ≥ 1 in control samples shown.

To further assess the noncanonical transcription and biogenesis of miR-4521, we examined a colon carcinoma cell line (RKO) that expresses high amounts of miR-4521 (but not miR-4638, which was not further examined). A variant of the cell line with doxycycline-inducible Cas9 (iCas9) was used for all experiments (de Almeida et al. 2021). To determine whether miR-4521 is dependent upon Pol III, RKO cells were treated with Pol III inhibitor ML- 60218, followed by sRNA-seq. miR-4521 was significantly depleted after 24 h of Pol III inhibition (Figure 7B). A few other miRNAs also showed Pol III-dependence; these include both strands of miR-1277 and guide strands miR-33a-5p, miR-153-3p, and miR-32-5p (Figure 7B, Table S13). Like miR-4521, the miR-33a-5p locus also exhibits Pol III ChIP (Ozsolak et al. 2008). Next, we examined DGCR8 and Dicer dependence by introducing dual guide RNAs (dgRNAs) targeting these loci (de Almeida et al. 2021). Cas9 expression was induced for four days, followed by harvest and western blot (which confirmed knockout of DGCR8 or Dicer, Figure S7E) and sRNA-seq. Knockout of DGCR8 or Dicer resulted in strong or moderate global depletion of miRNAs, respectively, when normalized to spike-ins (Figure 7C-D, Table S14-S15). Notably, miR-4521 expression was affected by neither DGCR8 nor Dicer knockout (Figure 7C- D). Thus, evolutionarily young small RNAs may also arise from Pol III transcription in humans.

## Discussion

In this study, we identified a new noncanonical miRNA biogenesis pathway that circumvents cleavage by the MP but relies on Dicer processing (Figures 1, 4). Using a variety of techniques, we have determined that the *mir-1829* family in *C. elegans* is derived from independent transcripts that are transcribed by Pol III (Figures 2, 3). Although these miRNA loci reside in the long introns of three Pol II-transcribed host genes, there is no detectable contribution of Pol II in *mir-1829* transcription (Figure 3F). Many classes of Pol III transcripts – including snaR-A, tRNAs, and SINEs – generate miRNAs via cleavage of a larger functional transcript (Babiarz et al. 2008; Martinez et al. 2017; Maute et al. 2013; Stribling et al. 2021). Unlike these transcripts, the *mir-1829* family exists exclusively as independent miRNA transcripts. The small RNAs generated by these loci are the only detectable species by northern blot, demonstrating that the *mir-1829* family members are the primary product of these loci, thus differentiating this unique biogenesis pathway from previously identified Pol III-transcribed miRNAs.

Given that the Pol III-transcribed *mir-1829* family resides in the long introns of its Pol II-transcribed host genes, we speculate that there may be competition for Pol II and Pol III for the F20D1.3*/ mir-1829b* locus. When the *mir-1829b* locus is expressed from an ectopic location in the *mir-1829 family(null)* background, *mir-1829b-5p* expression dramatically increases in comparison to levels in wild type (Figure 3C). Additionally, inhibiting F20D1.3 transcription by removing its promoter region – and thereby removing Pol II occupancy – results in a modest increase of *mir-1829b-5p* expression (Figure 3F). These results suggest that there is transcription interference of Pol III by Pol II at this locus. Previous studies have shown that Pol II can repress the transcription of tRNA genes (Lukoszek et al. 2013; Gerber et al. 2020). Since the *mir-1829* family is considered under the same class of Pol III transcripts as tRNAs (i.e. Type II), we hypothesize Pol II has an inhibitory effect on *mir-1829b* transcription. Additionally, it is a plausible that elongating Pol II physically occludes assembly of Pol III transcriptional machinery, limiting the transcriptional output of the *mir-1829* family (Martens et al. 2004; Petruk et al. 2006; Bird et al. 2006; Corbin and Maniatis 1989). However, the relationship between Pol II and Pol III is complex as emerging reports suggest that Pol II can promote Pol III transcription and vice versa (Gerber et al. 2020; Jiang et al. 2022; Listerman et al. 2007; K C et al. 2024). Pol II occupancy may also play a role in the differential spatial expression of the *mir- 1829* family and its host genes. This could be driven by chromatin remodeling during germ cell maturation, allowing for transcription of the miRNA but not the host gene in the distal germline (Yague-Sanz et al. 2023).

While we have demonstrated that the *mir-1829* family miRNAs require Pol III and Dicer for their biogenesis, additional required factors remain unidentified. We propose that a nuclear exonuclease may mature the 3’ end of the precursor by trimming this flank. Mutation of the proximal Pol III terminator reduces *mir-1829b-5p* production, suggesting the necessity of a very short flank 3’ of the precursor (Figure 3G); length heterogeneity at the 3’ end of 3p strands of *mir-1829* further suggest exonucleolytic trimming of the short trailing flank. The 3p strands also display some adenylated isoforms that are dependent on the enzyme GLDR-2. Our model (Figure 5E) is that the 3’ end of the miRNA precursor is matured and adenylated in the nucleus because 1) GLDR-2’s substrates show a strong 3p bias genome-wide beyond just the *mir-1829* family, suggesting its substrates are miRNA precursors (Vieux et al. 2021) and 2) GLDR-2 is localized to the nucleus (Figure 5D). Other interpretations in which GLDR-2 acts on the mature miRNA species are also possible but less parsimonious because they would require Dicer action in the nucleus or GLDR-2 activity in the cytoplasm. Importantly, GLDR-2 only modifies a small fraction of *mir-1829-3p* reads and is not required for biogenesis since its knockout does not perturb *mir-1829* family abundance (Figure 5C). If cleavage that generates the 5’ end of the precursor also occurs in the nucleus, then the precursor would be indistinguishable from a canonical miRNA precursor, enabling nuclear export. Spatio-temporal control of the unidentified nuclease(s) in *mir-1829* biogenesis could contribute to the localized enrichment of *mir-1829* in the distal germline.

Building on previous work that characterized the expression patterns and binding complements of *C. elegans* Argonaute proteins, we show that the *mir-1829* family’s expression is strongly germline enriched. Multiple lines of evidence suggest that *mir-1829b-5p* is very highly germline enriched, whereas a larger portion of *mir-1829c-5p* is expressed in the soma. An unusually large portion of the *mir-1829* family is loaded in the newly-characterized germline-specific Argonaute ALG-5. While two roles of ALG-5 in germline biology have been demonstrated, the targets of ALG-5 in these processes have yet to be discovered. Here we demonstrate that ALG-5 represses a tethered RNA (Figure 2); thus, ALG-5 likely to acts redundantly with ALG-1/2 on a shared set of target RNAs via their shared small RNA cargo. Given that ALG-5 functionally represses a tethered mRNA in the distal germline of adult hermaphrodites, which coincides with peak *mir-1829* expression, stability of the *mir-1829* family may be maintained within the spatial domain of ALG-5 activity. Furthermore, the rapid clearance of *mir-1829* in Dicer knockdown conditions suggests rapid turnover of *mir-1829* miRNAs that may determine or reinforce their spatial expression domain.

What is the contribution of the *mir-1829* family to animal physiology? Despite their enrichment in the germline, this miRNA family is not required for viability, germline function, or the robustness of the mitosis to meiosis transition. While these miRNAs are *C. elegans*- specific, they are preserved in the numerous wild isolates that have been collected around the world, suggesting that they may provide a selective advantage (Crombie et al. 2024). An interesting aspect of their future study will be to determine whether the 5p or 3p strands are the biologically important species. Although the 5p strands are more abundant, the 3p strands have greater sequence conservation, including across the functional seed region.

We expanded our search for additional noncanonical and Pol III-dependent miRNAs by comparing datasets in which MP, Dicer, or Pol III function are inactivated. In *C. elegans*, we identify 1) misannotated and unannotated 26G and 21U RNAs, 2) a pair of identical MP-independent small RNAs (*mir-8196a/b)* whose biogenesis may rely on their close proximity to a 21U RNA and 3) a Pol III-dependent small RNA (*mir-4937*) that requires neither MP nor Dicer for its biogenesis. In human cells, we identify multiple candidates for Pol III-transcribed small RNAs, including miR-32-5p, miR-33a-5p, miR-153-3p, miR-1277-3p, mir-1277-5p, and miR- 4521. miR-4521 is also DGCR8- and Dicer-independent. Whether miR-4521 functions as a miRNA is currently debated since it has shown activity in luciferase assays in some contexts, but also displays low Ago association in one study (Feng et al. 2019; Xing et al. 2021; Sun et al. 2021; Kuthethur et al. 2023; Orang et al. 2025). Notably, all of the newly identified noncanonical small RNAs are recently-evolved. Since miRNA loci are predicted to gradually evolve toward perfect biogenesis substrates (Berezikov 2011; Liu et al. 2008), these small RNAs and the *mir- 1829* family may be substrates for further evolution to canonical miRNAs.

De novo miRNAs are likely to target many mRNAs, having potentially deleterious effects and preventing their retention in the genome. Chen and Rajewsky suggested a model for miRNA gene evolution in which miRNAs first gain a low level of expression, possibly in a tissue-restricted manner (Chen and Rajewsky 2007). Then, deleterious target sites can be purged, and finally, miRNA level and domain of expression can increase, providing a selective advantage by repressing the remaining target mRNAs. The *mir-1829* family and other identified noncanonical small RNAs like *mir-8196a/b* and miR-4521 may be in the early stages of this trajectory, given their low level and tissue-restricted expression. Overall, these miRNAs fit multiple theoretical predictions of the characteristics of young de novo miRNA loci, thus representing a snapshot along the potential evolutionary trajectory to conserved canonical miRNA genes.

## Materials and methods

### C. elegans growth and maintenance

*C. elegans* were maintained on NGM plates seeded with OP50 at 20°C except where otherwise noted (Stiernagle 2006). For all experiments, embryos were harvested from gravid adults by hypochlorite treatment and washed thoroughly with M9 before hatching overnight in M9 supplemented with 1mM cholesterol at 20°C. Synchronized L1s were grown at 20°C to the appropriate stage as indicated. Strains and alleles used in this study are listed in Tables S1 and S2. For experiments using the auxin-inducible degron, synchronized L1s were grown for 48 h on NGM plates at 20°C, then washed onto 4mM auxin NGM plates at the L4 stage and grown for an additional 24 h at 20°C before harvesting.

### CRISPR/Cas9-mediated genome editing

All strains generated in this study were made by CRISPR-mediated editing as previously described (Yang et al. 2020). Deletions of the *mir-1829* family members and ALG-5 tethering reporter strains were generated by simultaneous CRISPR of two loci, in which one or both repair templates were integrated as indicated in Table S2.

### Brood size and embryonic viability assays

Brood size assays were conducted at 25°C as previously described (Kotagama et al. 2024). For *glp-1(ar202)* experiments, all strains were maintained at 15°C. A synchronized egg lay was performed for 3 h to isolate age-matched cohorts; these embryos were allowed to hatch at 15°C and then upshifted to 20°C 24 h after the synchronized egg lay. The worms were then kept at 20°C and single worms were followed to begin brood counts as above.

### Immunoprecipitation

Synchronized L1s were grown at 20°C for approximately 65 h (to first day of adulthood) prior to pellet collection. Frozen pellets were resuspended with volume of 2x Lysis buffer corresponding to the volume of the pellet. 1x Lysis buffer [30mM HEPES pH 7.4, 50mM KCl, 0.1% Triton X- 100, 2mM MgCl_2_, 10% glycerol, 2mM DTT, 80units/mL of RNAse inhibitor, 1x cOmplete Mini protease inhibitor cocktail (Roche)] was added to bring volume of all the samples to 1mL. Samples were subjected to 4-6 rounds of sonication (10 cycles of 30s on, 30s off per round) in a Bioruptor Pico, then clarified by centrifugation at >15,000*g* for 20 min at 4°C. Pierce Reducing Agent Compatible BCA Protein Assay was used to quantify protein prior to IP. IP inputs were 1ml of lysate at 5mg/mL for ALG-5 IP or 2mg/mL for all other samples. Inputs were combined with 50µl of washed Anti-FLAG M2 Magnetic Bead slurry (Millipore Sigma) and incubated with gentle agitation at 4°C overnight. After three washes (500µl of 30 mM HEPES pH 7.4, 100mM KCl, 0.1% Triton X-100, 2mM MgCl_2_, 10% glycerol, 2mM DTT, 80units/mL of RNAse inhibitor, 1x cOmplete Mini protease inhibitor), beads were resuspended in 1ml of 1x lysis buffer. Input, supernatant, and IP samples were saved in 50µl aliquots which were either run on 4–20% TGX gels (Bio-Rad) or used for RNA extraction (see below).

### RNA isolation

Frozen samples were resuspended with four pellet volumes of Trizol reagent (Fischer Scientific), then subjected to three freeze-thaw cycles on dry ice, followed by vortexing for 15 min at room temperature. RNA isolation was then performed according to the Trizol manufacturer’s instructions.

For IP-qPCR samples, extra care was taken to recover an equal proportion of RNA from each sample from IP workflows (input, supernatant, and IP). To this end, 82µl dH_2_O and 2.5µl yeast RNA (10mg/mL - Invitrogen) were added to 50µl aliquots of input, supernatant, or IP. 400µl Trizol LS (Thermo Fisher Scientific) was added to each sample, followed by 15 min of vortexing at room temperature. 106.25µl chloroform was then added to each sample prior to a 10-min centrifugation at >15,000*g*. A consistent volume (200µl) of the aqueous phase was transferred to a new tube and mixed with an equal volume phenol: chloroform: isoamyl alcohol (25:24:1, pH=4.5). After 10-min centrifugation at >15,000*g*, 100µl of the aqueous phase was transferred to a new tube, and RNA was precipitated with one volume (100µl) isopropanol overnight at −20°C. Pellets were washed with 75% ethanol and resuspended in 28µl dH_2_O.

### Rapid amplification of cDNA ends (RACE)

5’ ends of primary *mir-1829* family transcripts were determined using 5’/3’ RACE Kit, 2nd Generation Kit (Sigma-Aldrich). For cDNA synthesis, 1µl of ∼1µg/µl RNA sample was used. The amplification steps were carried out using Q5 High-Fidelity PCR Kit (NEB). For primers used, see Table S2. Purified PCR products were cloned into pCR™-Blunt II-TOPO vector using Zero Blunt® TOPO® PCR Cloning Kit (Invitrogen).

### Taqman miRNA qPCR

For each sample, 1.66µl of total RNA at 200ng/µl was used per 5µl reverse transcription reaction using the TaqMan MicroRNA Reverse Transcription kit (ThermoFisher). Completed RT reactions were diluted 1:4 with dH_2_O. Then, 1.33µl of diluted product was used in a 5µl qPCR reaction with Taqman miRNA probes and Taqman Universal Mastermix II with UNG (ThermoFisher) and were run in triplicate on the Applied Biosystems QuantStudio Pro 6.

### Reverse transcription-quantitative polymerase chain reaction (RT-qPCR) for mRNAs

Reverse transcription qPCR of mRNA was performed using KAPA SYBR® FAST One-Step qRT-PCR Master Mix (2X) Kit (Fisher Scientific) and gene-specific primers on the Applied Biosystems QuantStudio Pro 6. The primer sequences are listed in Table S2. For each reaction,

1.25µl of total RNA at 10ng/µl was used in a 5µl qPCR reaction, performing technical triplicate reactions.

### Immunoblotting

*C. elegans* protein samples were prepared and quantified as previously described (Kotagama et al. 2024). Samples were run on Mini-PROTEAN TGX gels (Bio-Rad). All blots were blocked in TBST containing 5% BSA (DCR-1::AID experiments) or 5% milk for 10 min to 1 h at room temperature. Membranes were probed overnight at 4°C with primary antibody, followed by F(ab’)2-Goat anti-Mouse IgG (H+L) Secondary Antibody, HRP (ThermoFisher) diluted 1:5000. Primary antibodies used were anti-FLAG M2 antibody (Sigma) 1:1000, anti-Dicer (Cell Signaling, 3363) 1:500, anti-DGCR8 (Genuinbiotech, 62316) 1:500, anti-Alpha Tubulin antibody DM1A (Abcam, ab7291) 1:5000 (AID experiments) or AA4.3 (Developmental Studies Hybridoma Bank) 1:5000 (IP experiments). ProtoGlow ECL (National Diagnostics) was used for signal detection of FLAG-tagged proteins, whereas Pierce ECL Western Blotting Substrate (Thermo Scientific) was used for signal detection of tubulin and human proteins. Signal was measured using Amersham Imager 680 (GE) and quantified using Image Studio Lite. For re-probing, membranes were stripped at 50°C for 45 min with stripping buffer (2% SDS, 62.5mM Tris HCl pH 6.8, 0.8% β-mercaptoethanol).

### Northern blot

Total RNA was separated on 15% urea gel (1.5mm) and transferred to Amersham Hybond- NX membranes. Membranes were crosslinked using 0.16M l-ethyl-3-(3-dimethylaminopropyl) carbodiimide (EDC) (Sigma) in 0.13M 1-methylimidazole, pH=8 at 60°C for 30 min as previously described (Pall et al. 2007). DNA oligos complementary to miRNAs of interest and U6 were 5’ end-labelled with ATP-γ-[^32^P] unless otherwise noted. For MP depletion experiments, 25µg of RNA was loaded and high stringency conditions were used. High stringency conditions consisted of probing overnight in hybridization buffer (7% SDS, 0.2M NaPO_4_ pH=7.2) at 42°C, followed by four washes for 10 min at 42°C (2x with 2x SSPE, 0.1% SDS, 1x with 1x SSPE, 0.1% SDS and 1x with 0.5x SSPE, 0.1% SDS) before exposing blots. For Dicer depletion experiments, 10µg RNA was loaded, and low stringency conditions were used. Low stringency conditions consisted of probing overnight in ExpressHyb Hybridization Solution (Takara Bio USA) at 32°C, followed by one wash with 2x SSC + 0.1% SDS at 32°C for 10 min and then one wash with 1x SSC + 0.1% SDS at 32°C for another 10 min before exposing blots. In between each probe, membranes were stripped by incubating in a solution of boiling 0.04% SDS at room temperature for 20 min, with shaking, for a total of three times.

### RNA Structure Predictions

Structure of the primary and precursor sequences of *mir-1829* family members were predicted using the minimum free energy (MFE) structure on RNAfold (Gruber et al. 2008; Lorenz et al. 2011). The Vienna format output for the primary structure was loaded into RNAcanvas (Johnson and Simon 2023) and the predicted substructure of the precursor was applied when this region differed in the primary transcript structure prediction.

### Small RNA-seq – C. elegans samples

Library preparation was performed using the NEBNext Small RNA Library Prep Set for Illumina as previously described (Donnelly et al. 2022). Pooled samples were sequenced on a NextSeq 2000. Raw sequence data is accessible under GEO accession number GSE282009. Briefly, sequence analysis was performed on the NIH High Performance Computing Cluster. Libraries were trimmed using Cutadapt 4.4 (Martin 2011), mapped using bowtie2 2.5.1 with settings ‘–no-unal –end-to-end –sensitive’ (Langmead and Salzberg 2012), and the resulting bam files were sorted and indexed using samtools 1.19 (Danecek et al. 2021). Reads were assigned to miRNAs using htseq 2.0.4 with settings ‘--mode union --nonunique fraction -a 0’ (Anders et al. 2015). Analysis of miRNA expression using DESeq2 was performed similarly to previously described (Kotagama et al. 2024), with the modification that size factors were calculated by dividing spike-in reads in each sample by the mean spike-in reads per sample. Published small RNA datasets were analyzed similarly, except that reads were trimmed and filtered for quality using Trimgalore 0.6.7 (DOI 10.5281/zenodo.5127898) with default settings.

### Small RNA-seq – cell culture samples

Small RNA libraries were prepped as described previously with minor modifications (Reichholf et al. 2019; Mandlbauer et al. 2024). See spike-in sequence and concentrations outlined in Table S2. Library preps were pooled then sequenced on NEBNext 2000 with 60 single end settings.

Libraries were adaptor trimmed then de-multiplexed with Cutadapt 5.0. UMIs were trimmed with UMI-tools 11.5 before mapping to Homo_sapiens_NCBI_GRCh38 genome using Bowtie2 2.5.3. The resulting bam files were sorted and indexed using samtools 1.21. Reads were assigned to miRNAs using htseq 2.04. Analysis of miRNA expression was performed with DEseq2 including spike-in size factors.

### Fluorescence Imaging

GLDR-2 adult worms raised at 15°C were mounted in 15μl of M9 with 5mM levamisole on a 2% agarose pad. ALG-5 tethering reporter strains were maintained at 20°C and mounted with M9 supplemented with 50mM sodium azide. Slides were cover slipped then imaged on a Nikon C2 confocal microscope using a 20x or 60x water immersion objective.

### Pol III ChIP analysis

*C. elegans* Pol III ChIP peaks were accessed at ENCODE (Accession file ENCFF924VCM) (Araya et al. 2014). For human ChIP data analysis, low quality bases and adaptors were trimmed from raw sequence reads with cutadapt v2.7 using 12 parameters -q 20 -a AGATCGGAAGAGC-minimum-length 25. Trimmed reads were aligned to the human GRCh38.p14 genome assembly using Bowtie2 v2.3.5 (Langmead and Salzberg 2012). Multimapping reads were removed with samtools v1.9 (Li et al. 2009) using the “view” subcommand and the additional argument “-q 20”, and duplicate reads were removed with the Picard v2.21.4 (http://broadinstitute.github.io/picard/index.html) Mark Duplicates tool. Peaks were called for each replicate individually using macs v2.2.7.1 (Zhang et al. 2008). Pooled peaks were called by providing multiple BAMs of all the replicates to macs2. miRNAs containing prominent Pol III peaks were called using bedtools intersect with the miRBase annotation file for the respective species (Quinlan and Hall 2010).

### Cell Culture

iCas9-RKO (sex unspecified; American Type Culture Collection, CRL-2577) cells were cultured in RPMI 1640 (Fischer Scientific) supplemented with 10% FBS (Takara Bio), 4mM L-glutamine (Thermo Fisher Scientific), 1mM sodium pyruvate (Sigma-Aldrich) and 100units/mL/100units/mL penicillin/streptomycin (Sigma-Aldrich) (de Almeida et al. 2021). Lenti-293T lentiviral packaging cells (female, Clontech, 632180) were cultured in Dulbecco’s modified Eagle’s medium (Sigma-Aldrich) supplemented with 10% FBS, 4mM L-glutamine, 1mM sodium pyruvate, and 100units/mL /100μg/mL penicillin/streptomycin. Cell lines were maintained at 37°C with 5% CO_2_ and routinely tested for mycoplasma contamination (InvivoGen).

### Lentivirus production and infection

Semiconfluent Lenti-X cells were transfected with lentiviral plasmids; pCMVR8.74 helper (Addgene, 22036), pCMV-VSV-G (Addgene, 8454), and Dicer or DGCR8-targeting dgRNAs (pLentiV1-Dual-CRISPR-hU6-mU6-EF1as-mCherry-P2A-Puro) using polyethylenimine (PEI) transfection (MW 25,000, Polysciences). After 4 days, virus containing supernatant was harvested then concentrated with Lenti-X Concentration (Takara Bio) using manufacturer instructions.

iCas9-RKO cells were transduced with Dicer or DGCR8 targeting dgRNAs (pLentiV1-Dual-CRISPR-hU6-mU6-EF1as-mCherry-P2A-Puro) at >/= 0.4 MOI. Cells containing Dicer or DGCR8 dgRNA were selected for by 2µg/mL Puromycin (Thermo Fisher Scientific).

### Polymerase III inhibitor and Dicer and DGCR8 knockout sample collection

Semiconfluent iCas9-RKO cells were treated with DMSO control or 50µM Polymerase III inhibitor (Millipore Sigma, CAS 577784-91-9). At 24 h, cells were washed once with 1x PBS then harvested in Trizol.

Semiconfluent Dicer or DGCR8 dgRNA containing iCas9-RKO cells were induced for Cas9 expression via 500ng/μL Doxycycline (Sigma Aldrich). After 4 days, cells were harvested for protein and RNA extraction.

For protein extraction, cell pellets were lysed on ice with NP40 lysis buffer (50mM Tris-Cl pH 7.4, 150mM NaCl, 1% NP40, 1mM EDTA, 1mM EGTA) supplemented with cOmplete Protease Inhibitor Cocktail (Millipore Sigma). After 30 min, lysates were centrifugated at 17,000*g* for 15 min. Supernatant was collected and mixed with ¼ volume 4x Laemmli buffer (Bio-Rad) with 50mM DTT (Bio-Rad). Protein lysates were boiled for 5 min at 95°C and placed in −20°C until western blotting.

For RNA extraction, cell pellets were re-suspended in Trizol (Fischer Scientific) then immediately frozen in dry ice and stored in −80°C until extraction. Once all samples were collected, RNA extraction was performed with manufacturer instructions with the modification of supplementing isopropanol and 80% ethanol with 0.2mM DTT. Finally, purified RNA was resuspended in water supplemented with 0.1mM DTT.

## Acknowledgments

We thank members of the McJunkin lab and Baltimore Worm Club (particularly Erik Andersen) for helpful discussions and WormBase. We are grateful to Jakub Zmajkovic and Johannes Zuber for sharing protocols, iCas9-RKO cells, DGCR8 and Dicer dgRNA vectors. Some of the strains used in this study were provided by the Caenorhabditis Genetics Center (CGC), which is funded by NIH Office of Research Infrastructure Programs (P40 OD010440). We thank Eleanor Maine and Tim Schedl for helpful suggestions. This work supported by the NIDDK IRP (ZIADK075147).

## Author Contributions

R.S. and K.M. conceived the study. R.S. and R.T. acquired, analyzed, and interpreted the data.

K.F.V. generated reagents and acquired data. D.H. and A.Z. generated reagents. R.S., G.Y., and K.M analyzed primary and published deep sequencing data. R.S. and K.M wrote the manuscript.

## References

Aalto AP, Nicastro IA, Broughton JP, Chipman LB, Schreiner WP, Chen JS, Pasquinelli AE. 2018. Opposing roles of microRNA Argonautes during Caenorhabditis elegans aging. PLoS Genet 14.

Alberti C, Manzenreither RA, Sowemimo I, Burkard TR, Wang J, Mahofsky K, Ameres SL, Cochella L. 2018. Cell-type specific sequencing of microRNAs from complex animal tissues. Nat Methods 15: 283–289.

Anders S, Pyl PT, Huber W. 2015. HTSeq--a Python framework to work with high-throughput sequencing data. Bioinformatics 31: 166–169.

Aoki ST, Lynch TR, Crittenden SL, Bingman CA, Wickens M, Kimble J. 2021. C. elegans germ granules require both assembly and localized regulators for mRNA repression. Nat Commun 12.

Araya CL, Kawli T, Kundaje A, Jiang L, Wu B, Vafeados D, Terrell R, Weissdepp P, Gevirtzman L, MacE D, et al. 2014. Regulatory analysis of the C. Elegans genome with spatiotemporal resolution. Nature 512: 400–405.

Auyeung VC, Ulitsky I, McGeary SE, Bartel DP. 2013. Beyond secondary structure: primary-sequence determinants license pri-miRNA hairpins for processing. Cell 152: 844–858.

Babiarz JE, Ruby JG, Wang Y, Bartel DP, Blelloch R. 2008. Mouse ES cells express endogenous shRNAs, siRNAs, and other Microprocessor-independent, Dicer-dependent small RNAs. Genes Dev 22: 2773–2785.

Behlke MA, Dames SA, McDonald WH, Gould KL, Devor EJ, Walder JA. 2000. Use of high specific activity StarFire oligonucleotide probes to visualize low-abundance pre-mRNA splicing intermediates in S. pombe. Biotechniques 29: 892–897.

Berezikov E. 2011. Evolution of microRNA diversity and regulation in animals. Nat Rev Genet 12: 846–860.

Berezikov E, Chung WJ, Willis J, Cuppen E, Lai EC. 2007. Mammalian mirtron genes. Mol Cell 28: 328–336.

Bird AJ, Gordon M, Eide DJ, Winge DR. 2006. Repression of ADH1 and ADH3 during zinc deficiency by Zap1-induced intergenic RNA transcripts. EMBO J 25: 5726.

Bortolamiol-Becet D, Hu F, Jee D, Wen J, Okamura K, Lin CJ, Ameres SL, Lai EC. 2015. Selective Suppression of the Splicing-Mediated MicroRNA Pathway by the Terminal Uridyltransferase Tailor. Mol Cell 59: 217–228.

Brenner JL, Jyo EM, Mohammad A, Fox P, Jones V, Mardis E, Schedl T, Maine EM. 2022. TRIM-NHL protein, NHL-2, modulates cell fate choices in the C. elegans germ line. Dev Biol 491: 43–55.

Brosnan CA, Palmer AJ, Zuryn S. 2021. Cell-type-specific profiling of loaded miRNAs from Caenorhabditis elegans reveals spatial and temporal flexibility in Argonaute loading. Nat Commun 12: 2194.

Brown KC, Svendsen JM, Tucci RM, Montgomery BE, Montgomery TA. 2017. ALG-5 is a miRNA-associated Argonaute required for proper developmental timing in the Caenorhabditis elegans germline. Nucleic Acids Res 45: 9093–9107.

Cheloufi S, Dos Santos CO, Chong MMW, Hannon GJ. 2010. A dicer-independent miRNA biogenesis pathway that requires Ago catalysis. Nature 465: 584–589.

Chen K, Rajewsky N. 2007. The evolution of gene regulation by transcription factors and microRNAs. Nat Rev Genet 8: 93–103.

Chung WJ, Agius P, Westholm JO, Chen M, Okamura K, Robine N, Leslie CS, Lai EC. 2011. Computational and experimental identification of mirtrons in Drosophila melanogaster and Caenorhabditis elegans. Genome Res 21: 286–300.

Cifuentes D, Xue H, Taylor DW, Patnode H, Mishima Y, Cheloufi S, Ma E, Mane S, Hannon GJ, Lawson ND, et al. 2010. A novel miRNA processing pathway independent of Dicer requires Argonaute2 catalytic activity. Science 328: 1694–1698.

Consalvo CD, Aderounmu AM, Donelick HM, Aruscavage PJ, Eckert DM, Shen PS, Bass BL. 2024. Caenorhabditis elegans Dicer acts with the RIG-I-like helicase DRH-1 and RDE-4 to cleave dsRNA. Elife 13.

Corbin V, Maniatis T. 1989. Role of transcriptional interference in the Drosophila melanogaster Adh promoter switch. Nature 337: 279–282.

Crombie TA, McKeown R, Moya ND, Evans KS, Widmayer SJ, LaGrassa V, Roman N, Tursunova O, Zhang G, Gibson SB, et al. 2024. CaeNDR, the Caenorhabditis Natural Diversity Resource. Nucleic Acids Res 52: D850–D858.

Danecek P, Bonfield JK, Liddle J, Marshall J, Ohan V, Pollard MO, Whitwham A, Keane T, McCarthy SA, Davies RM. 2021. Twelve years of SAMtools and BCFtools. Gigascience 10.

de Almeida M, Hinterndorfer M, Brunner H, Grishkovskaya I, Singh K, Schleiffer A, Jude J, Deswal S, Kalis R, Vunjak M, et al. 2021. AKIRIN2 controls the nuclear import of proteasomes in vertebrates. Nature 599: 491–496.

De Lencastre A, Pincus Z, Zhou K, Kato M, Lee SS, Slack FJ. 2010. MicroRNAs both promote and antagonize longevity in C. elegans. Curr Biol 20: 2159–2168.

Denli AM, Tops BBJ, Plasterk RHA, Ketting RF, Hannon GJ. 2004. Processing of primary microRNAs by the Microprocessor complex. Nature 432: 231–235.

Dexheimer PJ, Wang J, Cochella L. 2020. Two MicroRNAs Are Sufficient for Embryonic Patterning in C. elegans. Curr Biol 30: 5058–5065.e5.

Diag A, Schilling M, Klironomos F, Ayoub S, Rajewsky N. 2018. Spatiotemporal m(i)RNA Architecture and 3′ UTR Regulation in the C. elegans Germline. Dev Cell 47: 785–800.e8.

Dieci G, Fiorino G, Castelnuovo M, Teichmann M, Pagano A. 2007. The expanding RNA polymerase III transcriptome. Trends Genet 23: 614–622.

Donnelly BF, Yang B, Grimme AL, Vieux K-F, Liu C-Y, Zhou L, McJunkin K. 2022. The developmentally timed decay of an essential microRNA family is seed-sequence dependent. Cell Rep 40: 111154.

Duchaine TF, Wohlschlegel JA, Kennedy S, Bei Y, Conte D, Pang KM, Brownell DR, Harding S, Mitani S, Ruvkun G, et al. 2006. Functional Proteomics Reveals the Biochemical Niche of C. elegans DCR-1 in Multiple Small-RNA-Mediated Pathways. Cell 124: 343–354.

Dunham I, Kundaje A, Aldred SF, Collins PJ, Davis CA, Doyle F, Epstein CB, Frietze S, Harrow J, Kaul R, et al. 2012. An integrated encyclopedia of DNA elements in the human genome. Nature 489: 57–74.

Ender C, Krek A, Friedländer MR, Beitzinger M, Weinmann L, Chen W, Pfeffer S, Rajewsky N, Meister G. 2008. A human snoRNA with microRNA-like functions. Mol Cell 32: 519–528.

Fang W, Bartel DP. 2015. The Menu of Features that Define Primary MicroRNAs and Enable De Novo Design of MicroRNA Genes. Mol Cell 60: 131–145.

Feng X, Yan N, Sun W, Zheng S, Jiang S, Wang J, Guo C, Hao L, Tian Y, Liu S, et al. 2019. miR-4521-FAM129A axial regulation on ccRCC progression through TIMP-1/MMP2/MMP9 and MDM2/p53/Bcl2/Bax pathways. Cell Death Discov 5.

Fromm B, Domanska D, Høye E, Ovchinnikov V, Kang W, Aparicio-Puerta E, Johansen M, Flatmark K, Mathelier A, Hovig E, et al. 2020. MirGeneDB 2.0: the metazoan microRNA complement. Nucleic Acids Res 48: D132.

Grebert LFR, MacRae IJ. 2019. Regulation of microRNA function in animals. Nat Rev Mol Cell Biol 20: 21–37.

Gerber A, Ito K, Chu CS, Roeder RG. 2020. Gene-Specific Control of tRNA Expression by RNA Polymerase II. Mol Cell 78: 765–778.

Gregory RI, Yan KP, Amuthan G, Chendrimada T, Doratotaj B, Cooch N, Shiekhattar R. 2004. The Microprocessor complex mediates the genesis of microRNAs. Nature 432: 235–240.

Grishok A, Pasquinelli AE, Conte D, Li N, Parrish S, Ha I, Baillie DL, Fire A, Ruvkun G, Mello CC. 2001. Genes and mechanisms related to RNA interference regulate expression of the small temporal RNAs that control C. elegans developmental timing. Cell 106: 23–34.

Gruber AR, Lorenz R, Bernhart SH, Neuböck R, Hofacker IL. 2008. The Vienna RNA websuite. Nucleic Acids Res 36.

Gutiérrez-Pérez P, Santillán EM, Lendl T, Wang J, Schrempf A, Steinacker TL, Asparuhova M, Brandstetter M, Haselbach D, Cochella L. 2021. miR-1 sustains muscle physiology by controlling V-ATPase complex assembly. Sci Adv 7: eabh1434.

Ha M, Kim VN. 2014. Regulation of microRNA biogenesis. Nat Rev Mol Cell Biol 15: 509–524.

Han J, Lee Y, Yeom KH, Kim YK, Jin H, Kim VN. 2004. The Drosha-DGCR8 complex in primary microRNA processing. Genes Dev 18: 3016–3027.

Hansen TB, Venø MT, Jensen TI, Schaefer A, Damgaard CK, Kjems J. 2016. Argonaute-associated short introns are a novel class of gene regulators. Nat Commun 7: 11538.

Hussain S, Sajini AA, Blanco S, Dietmann S, Lombard P, Sugimoto Y, Paramor M, Gleeson JG, Odom DT, Ule J, et al. 2013. NSun2-mediated cytosine-5 methylation of vault noncoding RNA determines its processing into regulatory small RNAs. Cell Rep 4: 255–261.

Hutvágner G, McLachlan J, Pasquinelli AE, Bálint É, Tuschl T, Zamore PD. 2001. A cellular function for the RNA-interference enzyme Dicer in the maturation of the let-7 small temporal RNA. Science 293: 834–838.

Ikegami K, Lieb JD. 2013. Integral Nuclear Pore Proteins Bind to Pol III-Transcribed Genes and Are Required for Pol III Transcript Processing in C. elegans. Mol Cell 51: 840–849.

Jan CH, Friedman RC, Ruby JG, Bartel DP. 2011. Formation, Regulation and Evolution of Caenorhabditis elegans 3′UTRs. Nature 469: 97.

Jiang Y, Huang J, Tian K, Yi X, Zheng H, Zhu Y, Guo T, Ji X. 2022. Cross-regulome profiling of RNA polymerases highlights the regulatory role of polymerase III on mRNA transcription by maintaining local chromatin architecture. Genome Biol 23.

Jin W, Wang J, Liu C-P, Wang H-W, Xu R-M. 2020. Structural Basis for pri-miRNA Recognition by Drosha. Mol Cell 78: 423–433.e5.

Johnson PZ, Simon AE. 2023. RNAcanvas: interactive drawing and exploration of nucleic acid structures. Nucleic Acids Res 51: W501–W508.

K C R, Cheng R, Zhou S, Lizarazo S, Smith DJ, Van Bortle K. 2024. Evidence of RNA polymerase III recruitment and transcription at protein-coding gene promoters. Mol Cell 84: 4111–4124.e5.

Kang W, Fromm B, Houben AJ, Høye E, Bezdan D, Arnan C, Thrane K, Asp M, Johnson R, Biryukova I, et al. 2021. MapToCleave: High-throughput profiling of microRNA biogenesis in living cells. Cell Rep 37: 110015.

Kim K, Baek SC, Lee Y-Y, Bastiaanssen C, Kim J, Kim H, Kim VN. 2021. A quantitative map of human primary microRNA processing sites. Mol Cell 81: 3422–3439.e11.

Kim YK, Kim B, Kim VN. 2016. Re-evaluation of the roles of DROSHA, Export in 5, and DICER in microRNA biogenesis. Proc Natl Acad Sci U S A 113: E1881–E1889.

Knight SW, Bass BL. 2001. A role for the RNase III enzyme DCR-1 in RNA interference and germ line development in Caenorhabditis elegans. Science 293: 2269–2271.

Kotagama K, Grimme AL, Braviner L, Yang B, Sakhawala RM, Yu G, Benner LK, Joshua-Tor L, McJunkin K. 2024. Catalytic residues of microRNA Argonautes play a modest role in microRNA star strand destabilization in C. elegans. Nucleic Acids Res 52: 4985–5001.

Kozomara A, Birgaoanu M, Griffiths-Jones S. 2019. miRBase: from microRNA sequences to function. Nucleic Acids Res 47: D155–D162.

Kuscu C, Kumar P, Kiran M, Su Z, Malik A, Dutta A. 2018. tRNA fragments (tRFs) guide Ago to regulate gene expression post-transcriptionally in a Dicer-independent manner. RNA 24: 1093–1105.

Kuthethur R, Adiga D, Kandettu A, Jerome MS, Mallya S, Mumbrekar KD, Kabekkodu SP, Chakrabarty S. 2023. MiR-4521 perturbs FOXM1-mediated DNA damage response in breast cancer. Front Mol Biosci 10.

Langmead B, Salzberg SL. 2012. Fast gapped-read alignment with Bowtie 2. Nat Methods 9: 357–359.

Lee Y, Ahn C, Han J, Choi H, Kim J, Yim J, Lee J, Provost P, Rådmark O, Kim S, et al. 2003. The nuclear RNase III Drosha initiates microRNA processing. Nature 425: 415–419.

Lehrbach NJ, Castro C, Murfitt KJ, Abreu-Goodger C, Griffin JL, Miska EA. 2012. Post-developmental microRNA expression is required for normal physiology, and regulates aging in parallel to insulin/IGF-1 signaling in C. elegans. Rna 18: 2220–2235.

Lemus-Diaz N, Ferreira RR, Bohnsack KE, Gruber J, Bohnsack MT. 2020. The human box C/D snoRNA U3 is a miRNA source and miR-U3 regulates expression of sortin nexin 27. Nucleic Acids Res 48: 8074–8089.

Li H, Handsaker B, Wysoker A, Fennell T, Ruan J, Homer N, Marth G, Abecasis G, Durbin R. 2009. The Sequence Alignment/Map format and SAMtools. Bioinformatics 25: 2078–2079.

Liontis T, Verma K, Grishok A. 2023. DOT-1.1 (DOT1L) deficiency in C. elegans leads to small RNA-dependent gene activation. BBA advances 3.

Listerman I, Bledau AS, Grishina I, Neugebauer KM. 2007. Extragenic accumulation of RNA polymerase II enhances transcription by RNA polymerase III. PLoS Genet 3: 2268–2277.

Liu N, Okamura K, Tyler DM, Phillips MD, Chung WJ, Lai EC. 2008. The evolution and functional diversification of animal microRNA genes. Cell Res 18: 985–996.

Lorenz R, Bernhart SH, Höner zu Siederdissen C, Tafer H, Flamm C, Stadler PF, Hofacker IL. 2011. ViennaRNA Package 2.0. Algorithms Mol Biol 6.

Lu R, Yigit E, Li WX, Ding SW. 2009. An RIG-I-Like RNA Helicase Mediates Antiviral RNAi Downstream of Viral siRNA Biogenesis in Caenorhabditis elegans. PLoS Pathog 5: 1000286.

Lukoszek R, Mueller-Roeber B, Ignatova Z. 2013. Interplay between polymerase II- and polymerase III-assisted expression of overlapping genes. FEBS Lett 587: 3692–3695.

Mandlbauer A, Sun Q, Popitsch N, Schwickert T, Spanova M, Wang J, Ameres SL, Busslinger M, Cochella L. 2024. Mime-seq 2.0: a method to sequence microRNAs from specific mouse cell types. EMBO J 43: 2506.

Martens JA, Laprade L, Winston F. 2004. Intergenic transcription is required to repress the Saccharomyces cerevisiae SER3 gene. Nature 429: 571–574.

Martin M. 2011. Cutadapt removes adapter sequences from high-throughput sequencing reads. EMBnet J 17: 10–12.

Martinez G, Choudury SG, Slotkin RK. 2017. tRNA-derived small RNAs target transposable element transcripts. Nucleic Acids Res 45: 5142–5152.

Maute RL, Schneider C, Sumazin P, Holmes A, Califano A, Basso K, Dalla-Favera R. 2013. tRNA-derived microRNA modulates proliferation and the DNA damage response and is down-regulated in B cell lymphoma. Proc Natl Acad Sci U S A 110: 1404–1409.

Morgan PG, Sedensky MM. 1994. Mutations conferring new patterns of sensitivity to volatile anesthetics in Caenorhabditis elegans. Anesthesiology 81: 888–898.

Nguyen TL, Nguyen TD, Ngo MK, Nguyen TA. 2023. Dissection of the Caenorhabditis elegans Microprocessor. Nucleic Acids Res 51: 1512–1527.

Okamura K, Hagen JW, Duan H, Tyler DM, Lai EC. 2007. The mirtron pathway generates microRNA-class regulatory RNAs in Drosophila. Cell 130: 89–100.

Oler AJ, Alla RK, Roberts DN, Wong A, Hollenhorst PC, Chandler KJ, Cassiday PA, Nelson CA, Hagedorn CH, Graves BJ, et al. 2010. Human RNA polymerase III transcriptomes and relationships to Pol II promoter chromatin and enhancer-binding factors. Nat Struct Mol Biol 17: 620–628.

Orang A, Warnock NI, Migault M, Dredge BK, Bert AG, Bracken JM, Gregory PA, Pillman KA, Goodall GJ, Bracken CP. 2025. Chasing non-existent “microRNAs” in cancer. Oncogenesis 14.

Ozsolak F, Poling LL, Wang Z, Liu H, Liu XS, Roeder RG, Zhang X, Song JS, Fisher DE. 2008. Chromatin structure analyses identify miRNA promoters. Genes Dev 22: 3172–3183.

Pall GS, Codony-Servat C, Byrne J, Ritchie L, Hamilton A. 2007. Carbodiimide-mediated cross-linking of RNA to nylon membranes improves the detection of siRNA, miRNA and piRNA by northern blot. Nucleic Acids Res 35: e60.

Partin AC, Zhang K, Jeong B-C, Herrell E, Li S, Chiu W, Nam Y. 2020. Cryo-EM Structures of Human Drosha and DGCR8 in Complex with Primary MicroRNA. Mol Cell 78: 411–422.e4.

Persson H, Kvist A, Vallon-Christersson J, Medstrand P, Borg A, Rovira C. 2009. The non-coding RNA of the multidrug resistance-linked vault particle encodes multiple regulatory small RNAs. Nat Cell Biol 11: 1268–1271.

Petruk S, Sedkov Y, Riley KM, Hodgson J, Schweisguth F, Hirose S, Jaynes JB, Brock HW, Mazo A. 2006. Transcription of bxd Noncoding RNAs Promoted by Trithorax Represses Ubx in cis by Transcriptional Interference. Cell 127: 1209–1221.

Quinlan AR, Hall IM. 2010. BEDTools: a flexible suite of utilities for comparing genomic features. Bioinformatics 26: 841–842.

Reichholf B, Herzog VA, Fasching N, Manzenreither RA, Sowemimo I, Ameres SL. 2019. Time-resolved small RNA sequencing unravels the molecular principles of microRNA homeostasis. Mol Cell 75: 756.

Reimão-Pinto MM, Ignatova V, Burkard TR, Hung JH, Manzenreither RA, Sowemimo I, Herzog VA, Reichholf B, Fariña-Lopez S, Ameres SL. 2015. Uridylation of RNA Hairpins by Tailor Confines the Emergence of MicroRNAs in Drosophila. Mol Cell 59: 203–216.

Roberts JT, Cooper EA, Favreau CJ, Howell JS, Lane LG, Mills JE, Newman DC, Perry TJ, Russell ME, Wallace BM, et al. 2013. Continuing analysis of microRNA origins: Formation from transposable element insertions and noncoding RNA mutations. Mob Genet Elements 3: e27755.

Ruby JG, Jan CH, Bartel DP. 2007. Intronic microRNA precursors that bypass Drosha processing. Nature 448: 83–86.

Rybak-Wolf A, Jens M, Murakawa Y, Herzog M, Landthaler M, Rajewsky N. 2014. A variety of dicer substrates in human and C. elegans. Cell 159: 1153–1167.

Seroussi U, Lugowski A, Wadi L, Lao RX, Willis AR, Zhao W, Sundby AE, Charlesworth AG, Reinke AW, Claycomb JM. 2023. A comprehensive survey of c. Elegans argonaute proteins reveals organismwide gene regulatory networks and functions. Elife 12.

Sheng P, Fields C, Aadland K, Wei T, Kolaczkowski O, Gu T, Kolaczkowski B, Xie M. 2018. Dicer cleaves 5’-extended microRNA precursors originating from RNA polymerase II transcription start sites. Nucleic Acids Res 46: 5737–5752.

Simon DJ, Madison JM, Conery AL, Thompson-Peer KL, Soskis M, Ruvkun GB, Kaplan JM, Kim JK. 2008. The microRNA miR-1 regulates a MEF-2-dependent retrograde signal at neuromuscular junctions. Cell 133: 903–915.

Stiernagle T. 2006. Maintenance of C. elegans. WormBook 1–11.

Stribling D, Lei Y, Guardia CM, Li L, Fields CJ, Nowialis P, Opavsky R, Renne R, Xie M. 2021. A noncanonical microRNA derived from the snaR-A noncoding RNA targets a metastasis inhibitor. RNA 27: 694–709.

Stubna MW, Shukla A, Bartel DP. 2024. Widespread destabilization of Caenorhabditis elegans microRNAs by the E3 ubiquitin ligase EBAX-1. RNA 31: 51–66.

Stutzman A V., Liang AS, Beilinson V, Ikegami K. 2020. Transcription-independent TFIIIC-bound sites cluster near heterochromatin boundaries within lamina-associated domains in C. elegans. Epigenetics Chromatin 13: 1.

Sun B, Cong D, Chen K, Bai Y, Li J. 2021. Prognostic value of microRNA-4521 in non-small cell lung cancer and its regulatory effect on tumor progression. Open Medicine (Poland) 16: 1150–1159.

Tabara H, Yigit E, Siomi H, Mello CC. 2002. The dsRNA Binding Protein RDE-4 Interacts with RDE-1, DCR-1, and a DExH-Box Helicase to Direct RNAi in C. elegans. Cell 109: 861–871.

Vasquez-Rifo A, Jannot G, Armisen J, Labouesse M, Bukhari SIA, Rondeau EL, Miska EA, Simard MJ. 2012. Developmental characterization of the microRNA-specific C. elegans Argonautes alg-1 and alg-2. PLoS One 7.

Vieux KF, Prothro KP, Kelley LH, Palmer C, Maine EM, Veksler-Lublinsky I, McJunkin K. 2021. Screening by deep sequencing reveals mediators of microRNA tailing in C. elegans. Nucleic Acids Res 49: 11167–11180.

Westholm JO, Lai EC. 2011. Mirtrons: microRNA biogenesis via splicing. Biochimie 93: 1897– 1904.

Xie M, Li M, Vilborg A, Lee N, Shu M Di, Yartseva V, Šestan N, Steitz JA. 2013. Mammalian 5′-Capped MicroRNA Precursors that Generate a Single MicroRNA. Cell 155: 1568–1580.

Xiang S, Tian Z, Zheng W, Yang W, Du N, Gu Y, Yin J, Liu H, Jia X, Huang D, et al. 2021. Hypoxia downregulated miR-4521 suppresses gastric carcinoma progression through regulation of IGF2 and FOXM1. Mol Cancer 20.

Xu W, Liu J, Qi H, Si R, Zhao Z, Tao Z, Bai Y, Hu S, Sun X, Cong Y, et al. 2024. A lineage-resolved cartography of microRNA promoter activity in C. elegans empowers multidimensional developmental analysis. Nat Commun 15: 2783.

Yague-Sanz C, Migeot V, Larochelle M, Bachand F, Wéry M, Morillon A, Hermand D. 2023. Chromatin remodeling by Pol II primes efficient Pol III transcription. Nature Communications 2023 14:1 14: 1–13.

Yang B, Schwartz M, McJunkin K. 2020. In vivo CRISPR screening for phenotypic targets of the mir-35-42 family in C. elegans. Genes Dev 34: 1227–1238.

Yang S, Maurin T, Robine N, Rasmussen KD, Jeffrey KL, Chandwani R, Papapetrou EP, Sadelain M, O’Carroll D, Lai EC. 2010. Conserved vertebrate mir-451 provides a platform for Dicer-independent, Ago2-mediated microRNA biogenesis. Proc Natl Acad Sci U S A 107: 15163–15168.

Zamudio JR, Kelly TJ, Sharp PA. 2014. Argonaute-bound small RNAs from promoter-proximal RNA polymerase II. Cell 156: 920–934.

Zhang Y, Liu T, Meyer CA, Eeckhoute J, Johnson DS, Bernstein BE, Nussbaum C, Myers RM, Brown M, Li W, et al. 2008. Model-based analysis of ChIP-Seq (MACS). Genome Biol 9.

