## Supplemental Figures for "A noncanonical Pol III-dependent, Microprocessor-independent biogenesis pathway generates a germline enriched miRNA family"

**List of Supplemental Figures and Tables**

**Figure S1.** Data related to Microprocessor-independence of *mir-1829.*

**Figure S2.** The germline-enriched *mir-1829* family is not required for fertility or viability.

**Figure S3.** Knock-in of *mir-1829b* transcriptional unit at heterologous locus generates mature *mir-1829b.*

**Figure S4.** The AID system allows for efficient depletion of DCR-1, RPC-1, and NSUN-2.

**Figure S5.** The *mir-1829* family is significantly reduced upon germline-specific Dicer depletion.

**Figure S6.** DRH-1, RDE-4, NSUN-2, and key cellular nucleases are dispensable for *mir-1829b/c* biogenesis.

**Figure S7.** Noncanonical miRNA biogenesis extends beyond the *mir-1829* family.

**Table S1.** Strains used in this study.

**Table S2.** Alleles generated and oligonucleotides used in this study.

**Table S3.** 5' RACE fragment positions relative to precursor.

**Table S4.** Samples used for deep sequencing.

**Table S5.** Raw miRNA and spike-in reads for all samples.

**Table S6.** Results of small RNA sequencing of Dicer knockdown in whole worms (soma & germline).

**Table S7.** Results of small RNA sequencing of Dicer knockdown in the germline.

**Table S8.** Expanded results of small RNA sequencing of annotated miRBase miRNAs in Dicer knockdown (soma & germline).

**Table S9.** Results of small RNA sequencing of RPC-1 knockdown in whole worms (soma & germline).

**Table S10.** Identification of additional noncanonical microRNAs.

**Table S11.** Summary of mis-annotated and noncanonical microRNAs.

**Table S12.** Expanded results of small RNA sequencing analysis of annotated miRBase miRNAs in germline-specific Dicer knockdown.

**Table S13.** Results of small RNA sequencing upon Pol III inhibition in iCas9-RKO cells.

**Table S14.** Results of small RNA sequencing after DGCR8 knockout in iCas9-RKO cells.

**Table S15.** Results of small RNA sequencing after DICER knockout in iCas9-RKO cells.

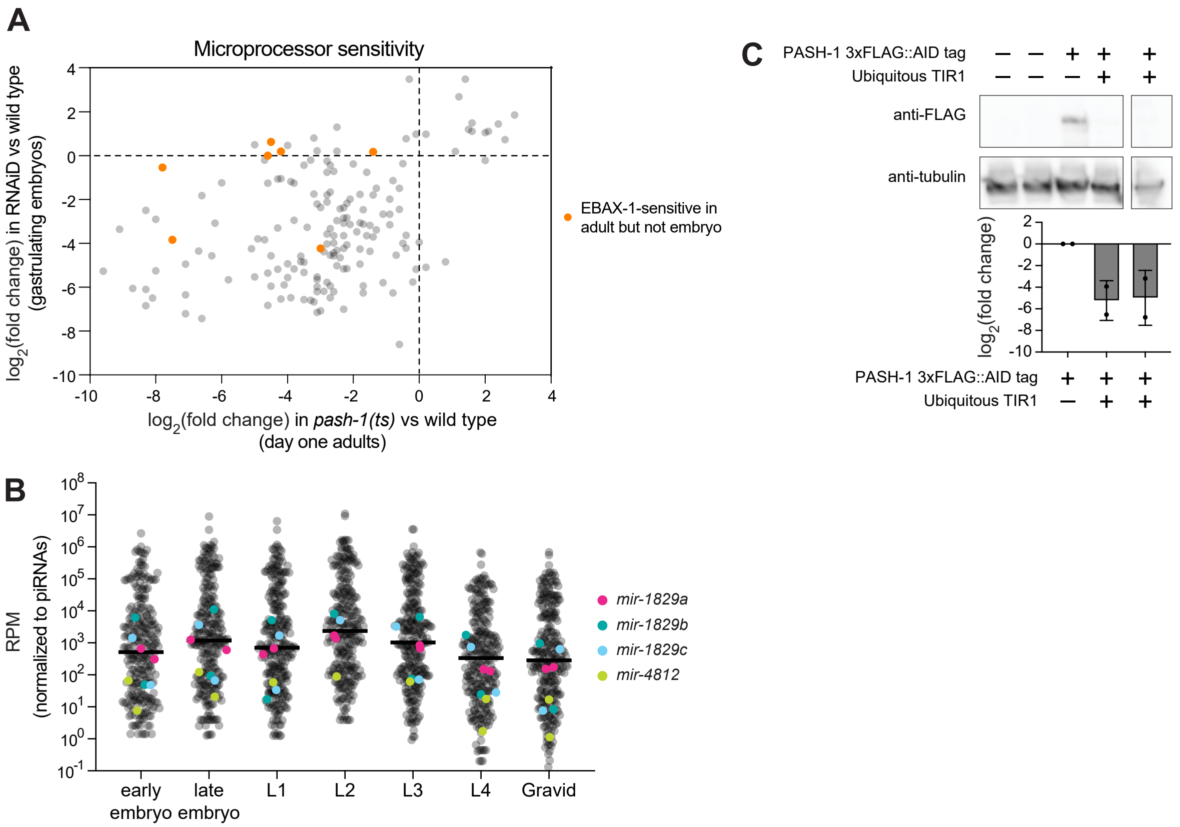


**Figure S1. Data related to Microprocessor-independence of *mir-1829*.** A) Comparison of log2(fold change) of miRNAs 24h post-upshift to restrictive temperature in *pash-1(ts)* vs. wild type (Vieux et al. 2021) (x-axis) to those in RNAiD depletion of the MP (Dexheimer et al. 2020) (y-axis) with highlighting of miRNAs that are stabilized in *ebax-1*$^{-/-}$ mutants in adults but not embryos. B) Relative abundance of *mir-1829* in relation to all miRNAs throughout *C. elegans* development from bias-minimized small RNA cloning from Stubna, et al. 2024. C) Representative western blot of PASH-1 knockdown using the AID system. Quantification of FLAG tag signal, normalized to tubulin signal, shown below. PASH-1 levels are normalized to PASH-1 AID tag without TIR1 (MLC1061) treated with auxin. Mean and SD shown.

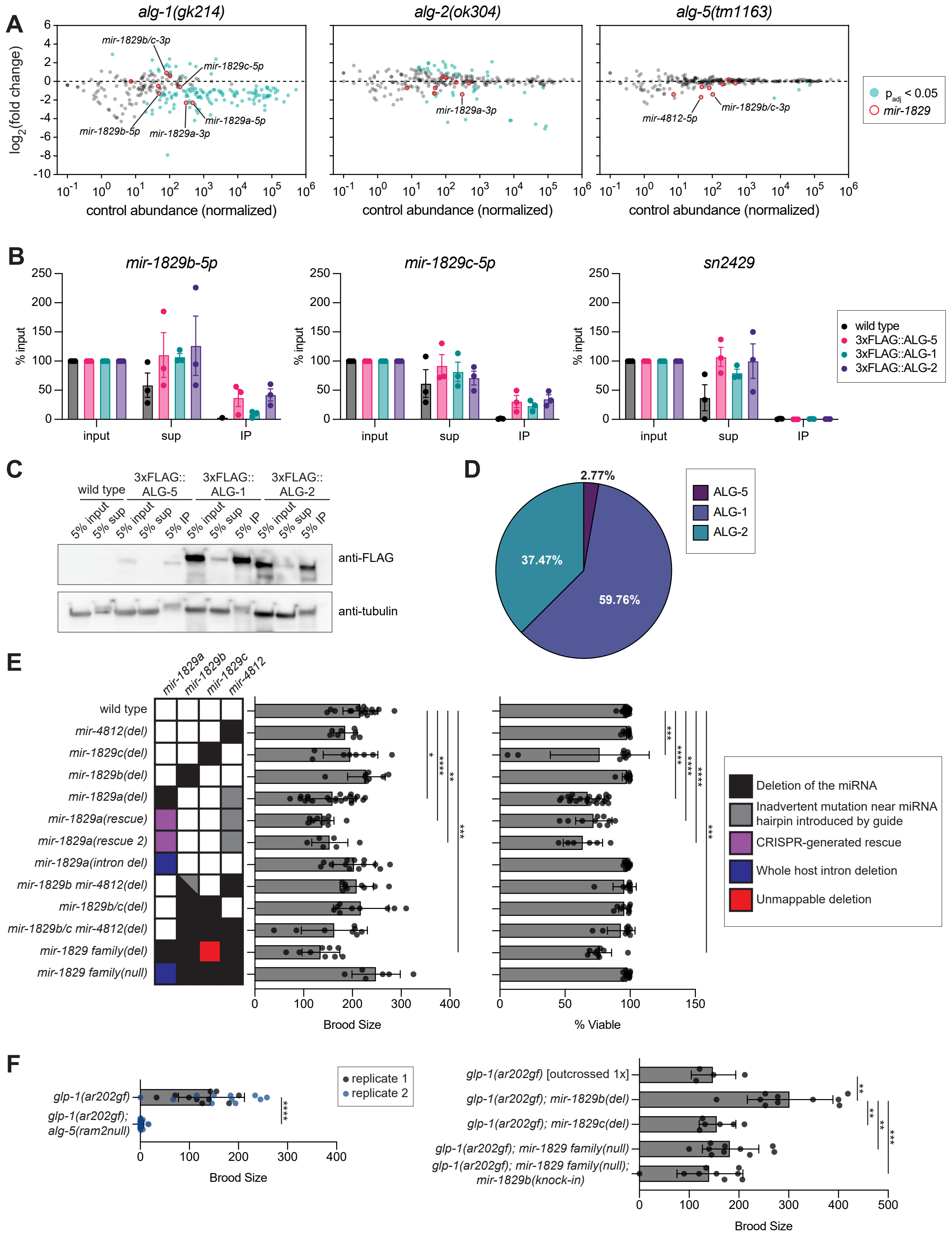


**Figure S2. The germline-enriched *mir-1829* family is not required for fertility or viability.** A) MA plots showing abundance in wild type on x-axis and $\log_2$(fold change) in Ago knockouts (*alg-1(gk214), alg-2(ok304), and alg-5(tm1163),* respectively) compared to wild type (N2) on y-axis from Seroussi, et al. 2023. miRNAs showing significant sensitivity to Ago mutants are shown in blue-green, and the *mir-1829* family is highlighted in red open circles. Three biological replicates of each genotype were analyzed using DESeq2. B) RT-qPCR of *mir-1829b/c* levels in input, supernatant (sup), and immunoprecipitations (IP) of miRNA Argonautes ALG-1, ALG-2, and ALG-5. Wild type lysate and non-miRNA sn2429 were included as controls. *mir-1829b-5p* is evenly distributed between ALG-2 and ALG-5, and *mir-1829c-5p* is equally loaded into all three miRNA Argonautes. Mean and SEM shown. C) Representative western blot of miRNA Argonaute IPs used for RT-qPCR. 5% of input, supernatant (sup) and IP was loaded into each well. D) Representation of relative miRNA Argonaute abundances in whole animal adult samples. E) CRISPR-generated mutants of the *mir-1829* family were assessed for fertility (left) and embryonic viability (right) defects at 25°C. F) Consistent with Brenner *et al.*, *glp-1(ar202); alg-5(ram2)* double mutant hermaphrodites produce no progeny when upshifted from 15°C to 20°C (left). Mutations of the *mir-1829* family members do not phenocopy *alg-5(ram2)* in *glp-1(ar202)* background (right). (E-F) Mean and SD shown. Two-way ANOVA. Two-way ANOVA. ****$p$<0.0001, ***$p$<0.001, **$p$<0.01 *$p$<0.05.

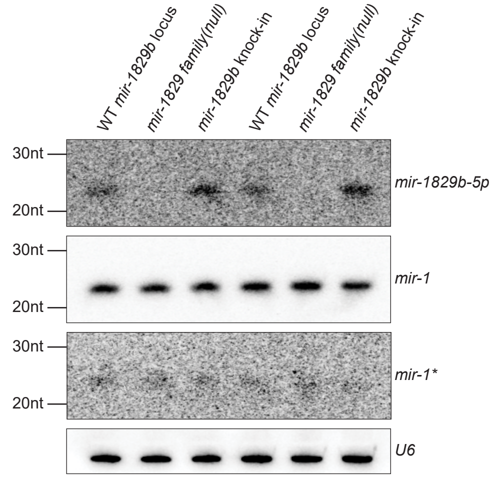


**Figure S3. Knock-in of *mir-1829b* transcriptional unit at heterologous locus generates mature *mir-1829b*.** Northern blot using low stringency conditions (see methods) demonstrates that re-integration of a 510bp minimal *mir-1829b* transcriptional unit in a *mir-1829 family(null)* mutant background expresses a mature small RNA (two biological replicates shown)

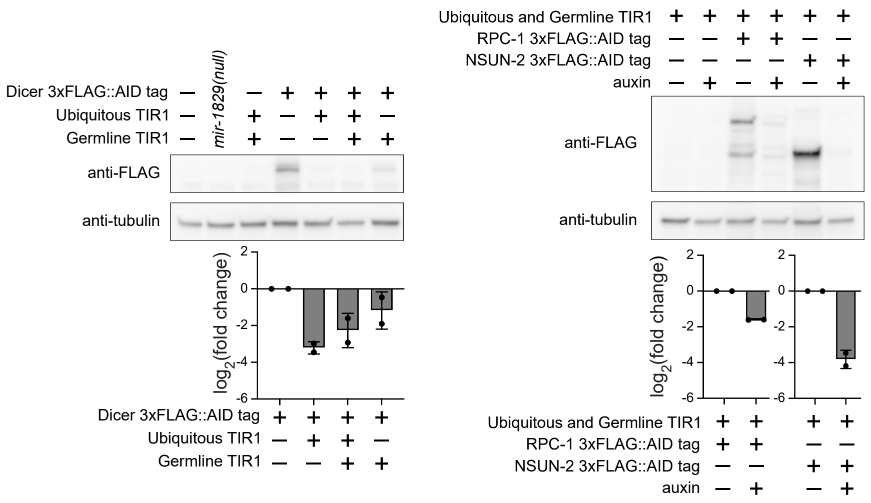


**Figure S4. The AID system allows for efficient depletion of DCR-1, RPC-1, and NSUN-2.** Representative westerns of Dicer/DCR-1 (left), RPC-1 and NSUN-2 knockdown (right) using the AID system. Quantification of FLAG tag signal normalized to tubulin signal shown below. Dicer levels are normalized to Dicer AID tag without TIR1 (UY212), and RPC-1 and NSUN-2 levels are normalized to samples not treated with auxin. Mean and SD shown.

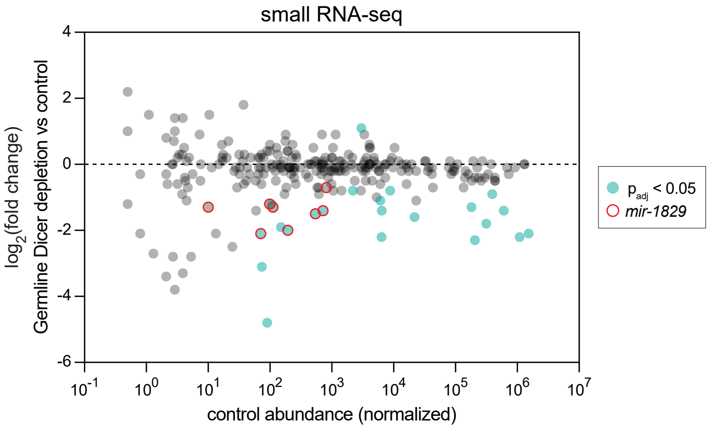


**Figure S5. The *mir-1829* family is significantly reduced upon germline-specific Dicer depletion.** Small RNA-seq. MA plot showing abundance in control (DCR-1::AID tag alone, UY212) on x-axis and $log_2$(fold change) in DCR-1::AID; germline TIR1 (MCJ660) compared to UY212 on y-axis. miRNAs showing significant sensitivity to Dicer depletion are shown in blue-green, and the *mir-1829* family is highlighted in red open circles. Three biological replicates of each genotype were analyzed using DESeq2.

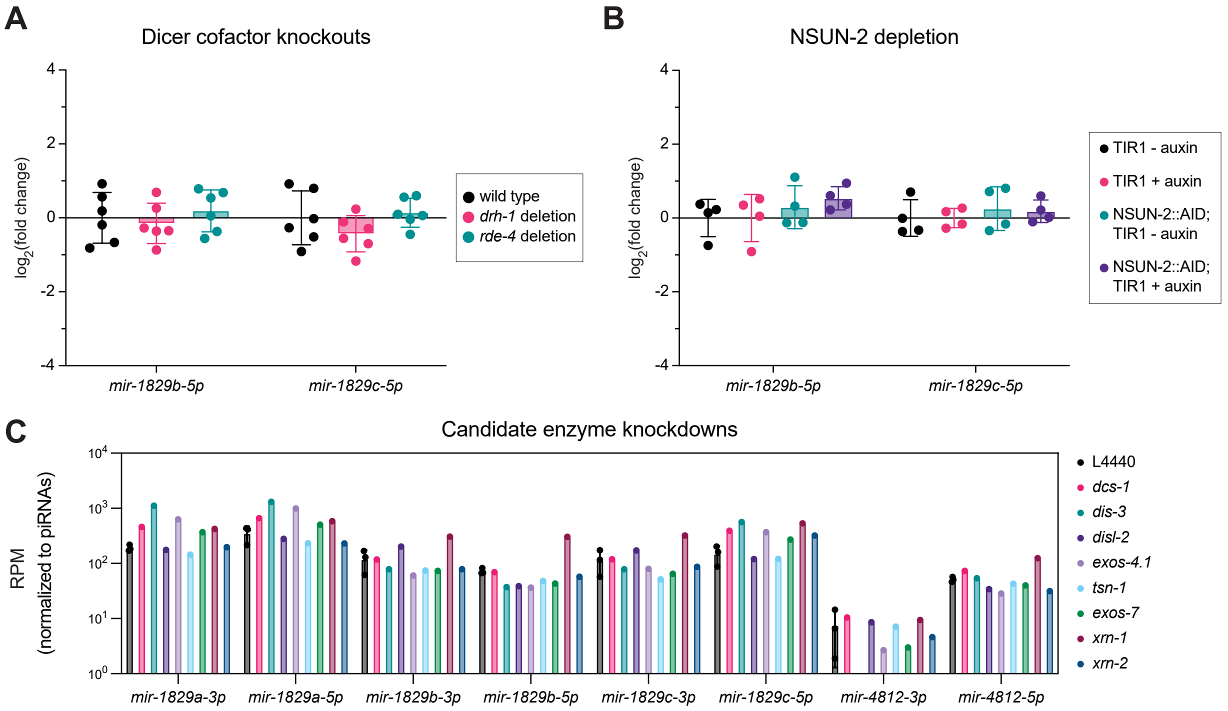


**Figure S6. DRH-1, RDE-4, NSUN-2, and key cellular nucleases are dispensable for *mir-1829b/c* biogenesis.** A) miRNA qPCR normalized to a small RNA control (*sn2429)* and further normalized to wild type levels. B) qPCR of miRNAs in AID-tagged NSUN-2 strain with ubiquitous TIR1 (MCJ677) or TIR1 alone (MLC1040). miRNA qPCR normalized to a small RNA control (*sn2429)*. MCJ677 without auxin treatment was further normalized to MLC1040 without auxin treatment, and auxin-treated MCJ677 was normalized to auxin-treated MLC1040. (A-B) Mean and SD shown. Two-way ANOVA showed no significant changes. C) Reads per million piRNA-mapping reads in adult samples grown on empty vector (L4440) or the indicated RNAi vector. Data was reanalyzed from Vieux et al. 2021.

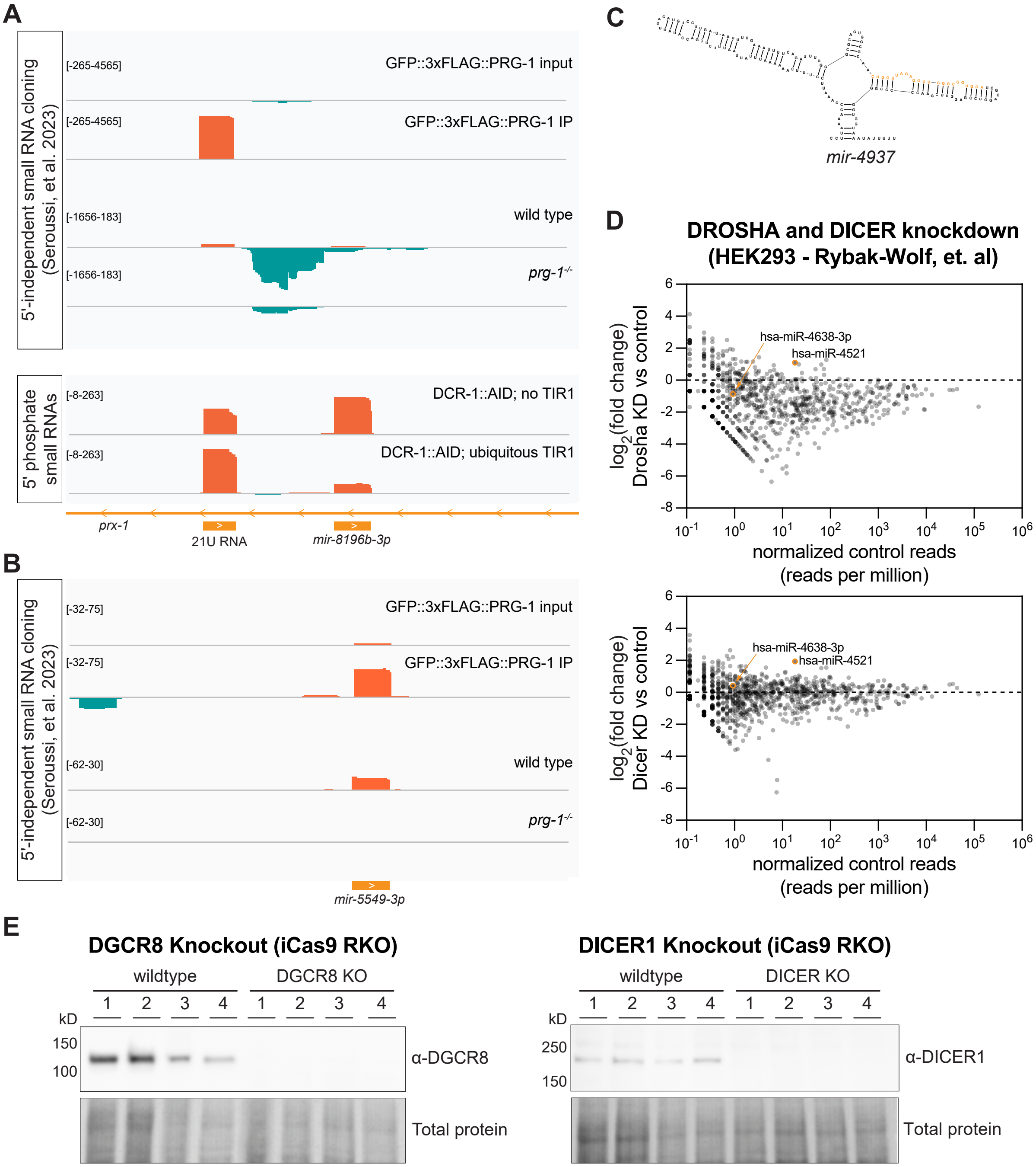


**Figure S7. Noncanonical miRNA biogenesis extends beyond the *mir-1829* family.** A) Genome browser tracks showing proximity of *mir-8196b* to an upstream recently-annotated 21U RNA. 5′-independent small RNA cloning following polyphosphatase treatment also shows 22G RNAs generated from the locus. *mir-8196b* but not the 21U shows Dicer-dependence. B) Genome browser tracks showing misannotated 21U RNA *mir-5549-3p,* which is enriched in PRG-1 IP and depleted in *prg-1*$^{-/-}$ mutants. C) Predicted secondary structure of *pri-mir-4937.* D) MA plot of small RNA sequencing from HEK293 cells in which DROSHA or DICER is knocked down by siRNA. *mir-4521* shows DROSHA- and DICER-independence (Rybak-Wolf, et al. 2014). E) Western blot demonstrating efficient knockout of DGCR8 or DICER1 in iCas9 RKO cells.
